# Dynamics of the DYNLL1/MRE11 complex regulates DNA end resection and recruitment of the Shieldin complex to DSBs

**DOI:** 10.1101/2023.03.27.534416

**Authors:** Rui Zhou, Michelle L. Swift, Aleem Syed, Kaimeng Huang, Lisa Moreau, John A. Tainer, Panagiotis A. Konstantinopoulos, Alan D. D’Andrea, Yizhou Joseph He, Dipanjan Chowdhury

## Abstract

Extent and efficacy of DNA end resection at DNA double strand break (DSB)s determines the choice of repair pathway. Here we describe how the 53BP1 associated protein DYNLL1 works in tandem with Shieldin and the CST complex to protect DNA ends. DYNLL1 is recruited to DSBs by 53BP1 where it limits end resection by binding and disrupting the MRE11 dimer. The Shieldin complex is recruited to a fraction of 53BP1-positive DSBs hours after DYNLL1 predominantly in the G1 cells. Shieldin localization to DSBs is dependent on MRE11 activity and is regulated by the interaction of DYNLL1 with MRE11. BRCA1-deficient cells rendered resistant to PARP inhibitors by the loss of Shieldin proteins can be re-sensitized by the constitutive association of DYNLL1 with MRE11. These results define the temporal and functional dynamics of the 53BP1-centric DNA end resection factors in cells.

## MAIN

Resection of a broken DNA end plays a determinant role in the choice of DNA double strand break (DSB) repair pathways. Homologous recombination (HR) mediated DSB repair occurs in the S phase and requires extended DNA end resection promoted by the tumor suppressor BRCA1 ^1^. Minimal end resection facilitates non-homologous end joining (NHEJ), an efficient pathway which remains active throughout the cell cycle^2, 3^. Extensive work in the past two decades has defined the end resection machinery, specifically the large number of factors that process DNA ends, such as the exo- and endonucleases and the helicases that function therein^4^. In the last decade, 53BP1 has emerged as a focal anti-resection factor that counters BRCA1 to favor NHEJ^5–10^. More recently, factors that functionally and biochemically associate with 53BP1 to restrict end resection have been discovered in genetic and biochemical screens^11^. These include the Shieldin complex^12–14^, composed of REV7, a known 53BP1-pathway component, and three uncharacterized proteins: C20orf196 (SHLD1), FAM35A (SHLD2) and CTC-534A2.2 (SHLD3). Shieldin localizes to DSB sites in a 53BP1-dependent manner, and directly binds to single-stranded DNA via the OB-fold domains of the SHLD2 subunit to protect DNA ends from hyper-resection^12^. Next the CST complex (CTC1-STN1-TEN1) interacts with Shieldin components and localizes with Polα to sites of DNA damage in a 53BP1- and Shieldin-dependent manner to fill in resected DSBs and counteract end-resection^7, 10^. Depleting components of either the Shieldin or CST complexes promote resection and HR, thereby suppressing the sensitivity of BRCA1-null cells to PARP inhibitors (PARPi)^7, 10^. The spatiotemporal dynamics of how these repair proteins function together to inhibit end-resection remains to be determined.

We discovered the cytoplasmic motor protein, DYNLL1 was an anti-resection factor that was recruited to DSBs and directly bound the nuclease MRE11 to impede DNA end resection *in vitro* and in cells^15^. Very similar to the Shieldin and CST complex, loss of DYNLL1 conferred resistance to PARPi sensitivty in BRCA1-mutant cell lines. Several key questions emerged from this work: (i) How and when is DYNLL1 recruited to DSBs? (ii) How does DYNLL impede the activity of MRE11? (iii) What is the relative order of recruitment of DYNLL1 and the Shieldin/CST complex? (iv) Most importantly, why does the cell utilize multiple independent complexes for its anti-resection activity at DNA ends? Here we address these questions and define the dynamics of these end resecting factors and how they regulate DSB repair. We observed that DYNLL1 recruitment to DSBs was dependent solely on 53BP1 and no other known 53BP1 associated anti-resection protein. This is distinct from the Shieldin and CST complexes which required REV7 and RIF1 proteins^7, 12^. DYNLL1 localization at DNA lesions occurred concurrently with 53BP1 to block the initiation of DNA end resection. Biochemical mapping of the interaction domain of MRE11 with DYNLL1 combined with *in vitro* studies revealed that DYNLL1 interferes with the dimerization of MRE11 and potentially removes it from chromatin. Shieldin and CST complex primarily localized to a subset of 53BP1 positive DSBs primarily in the G1 phase hours after DNA damage. MRE11 and CtIP were necessary for Shieldin localization. Loss of DYNLL1 promoted Shieldin/CST recruitment to DSBs. Conversely, constitutive interaction of DYNLL1 with MRE11 impaired Shieldin/CST recruitment to DSBs. Finally PARPi resistance induced in BRCA1-mutant cells by depletion of Shieldin/CST complex proteins was ‘rescued’ by the expression of a phosphomimetic DYNLL1 mutant that constitutively binds MRE11. Together our results establish DYNLL1 as a partner of 53BP1 in the anti-resection machinery that functions upstream of other 53BP1 interacting proteins to prevent DNA end resection.

## RESULTS

### DYNLL1 recruitment to DSBs is dependent on 53BP1 but independent of other 53BP1-associated factors

We observed that DYNLL1 is recruited to DSBs^15^. Using EGFP-tagged DYNLL1 and laser microirradiation-induced DSBs, we determined that DYNLL1 localized to DNA lesions rapidly within minutes (Fig. 1A), with kinetics comparable to 53BP1 (Extended Fig. 1A). This is consistent with observations that DYNLL1 directly interacts with 53BP1 near the oligomerization domain (1113aa-1177aa)^16–18^. Therefore, the constitutive interaction with 53BP1 allows DYNLL1 to be recruited with 53BP1 to DSBs. Loss of 53BP1 strongly impaired recruitment of DYNLL1 to laser micro-irradiated or IR induced DSBs (Extended Fig. 1B). Consistent with prior data, loss of 53BP1 also decreased DYNLL1 chromatin binding^17^ both in the presence and absence of damage (Extended Fig. 1C). Alanine substitution of the three anchor residues (GIQ and TQT) in both the DYNLL1-binding motifs of 53BP1 blocks oligomerization domain (OD)-independent 53BP1 localization^18^, thereby inhibiting DYNLL1 localization to chromatin^17^ and DYNLL1 recruitment to DSBs (Fig. 1B). This 53BP1 mutant localizes to DSBs and forms foci but does not recruit DYNLL1(Fig. 1B). It is noteworthy that BRCA1 deficiency had no impact on DYNLL1 foci (Fig. 1C and Extended Fig.1D-1F). We also confirmed previous findings that loss of DYNLL1 alone did not significantly impact recruitment of 53BP1 to DSBs^15, 18^, (Extended Data Fig. 1G). The other 53BP1 interactors, RIF1/PTIP/REV7 associate with 53BP1 after DNA damage induced ATM-mediated phosphorylation of the N-terminus^11^. Along with 53BP1, RIF1 and REV7 are necessary for recruitment of Shieldin complex to DSBs^19 7, 10, 12^. The Shieldin complex, specifically SHLD1, is required for the CST complex to be localized at DSBs^10, 20^. Therefore, we asked whether any of these proteins play a role in the recruitment or retention of DYNLL1 to DSBs. While recruitment of DYNLL1 to DSBs was dependent on 53BP1, depletion of 53BP1 associated factors, RIF1, REV7, and components of the Shieldin and CST complexes did not impact DYNLL1 localization to DSBs (Fig. 1E and Extended Data Fig. 1H). From these results we infer that recruitment of DYNLL1 to DSBs is solely dependent on 53BP1, and not the associated proteins.

**Fig. 1:**
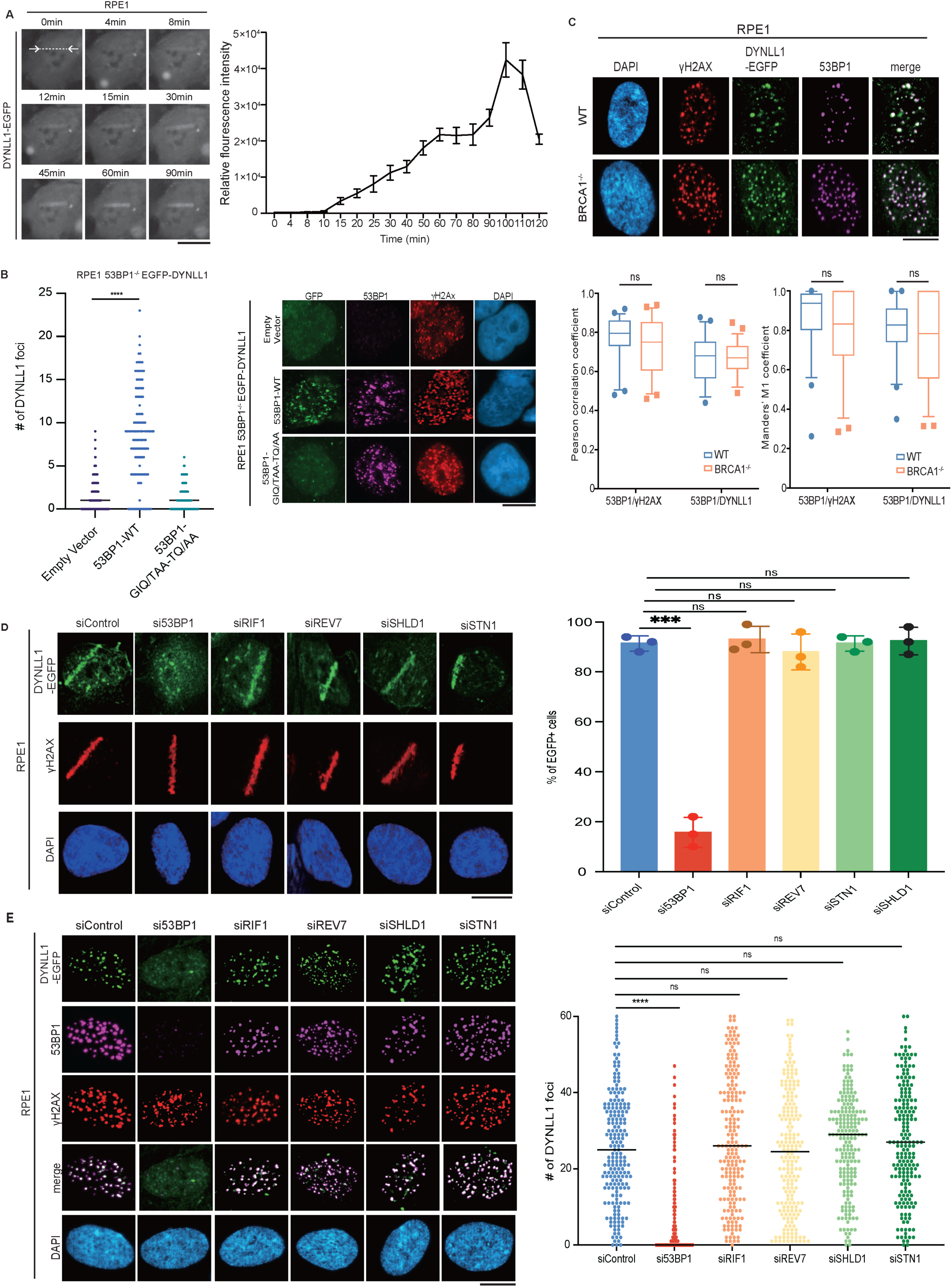
DYNLL1 recruitment to DSBs is dependent on 53BP1 but independent of other 53BP1-associated factors. (A) Live cell representative images of RPE1 cells expressing DYNLL1-EGFP after laser microirradiation. (B) RPE1 53BP1^-/-^ cells coexpressing EGFP-DYNLL1 and 53BP1-DYNLL1 binding mutants were subjected to 2 Gy irradiation and 2 hours later were processed for immunofluorescence using antibodies against GFP, 53BP1, and γH2Ax. (C) Representative immunofluorescent images of RPE1 wild-type or BRCA1^-/-^ cells 2 hours after exposure to 2 Gy irradiation. Cells were fixed and processed for immunofluorescence using antibodies against 53BP1, GFP (DYNLL1), and γH2Ax. (D, E) Representative images of RPE1 cells coexpressing GFP-tagged DYNLL1 and various siRNA constructs and subjected to laser-microirradiation (D) or 2Gy irradiation (E), fixed 1 hour later, and probed with antibodies against 53BP1, GFP (DYNLL1), and γH2Ax. (A-E) For all experiments, n=3 biologically independent experiments, counting ≥ 100 cells per experiment. P- values for foci quantification were calculated using the Mann-Whitney test. P-values for foci colocalization and “laser positive” cell analysis was calculated using unpaired t-tests. Bars on scatter plots represent the median. Error bars represent the mean±s.e.m.

### DYNLL1 regulates MRE11 activity independent of 53BP1

53BP1 influences DSB repair pathway choice and PARPi sensitivity by regulating DNA end resection at DSBs. In response to DNA damage DYNLL1 loss increased MRE11 foci formation in BRCA1 deficient cells ^15^ (Fig. 2A). Consistent with its impact on DYNLL1 foci formation, loss of 53BP1 also increased MRE11 foci formation in BRCA1-deficient/mutant cells (Fig. 2A). Intriguingly, BRCA1 loss diminished MRE11 foci and this was ‘rescued’ by the co-deletion of 53BP1 (Fig. 2B). Next we depleted MRE11 to investigate it’s importance in the BRCA1/53BP1/DYNLL1 regulatory loop in the context of PARPi treatment. PARPi sensitivity in BRCA1^-/-^ (or BRCA1 mutant) 53BP1^-/-^ was rescued upon depletion of MRE11 (Fig. 2C and Extended Fig. 2A,B). This suggested that MRE11 functioned downstream of 53BP1 in regulating DNA end resection. Importantly, depletion of MRE11 further sensitized BRCA1^-/-^ cells to PARPi (Fig. 2C). The implication was that the impact of MRE11 on PARPi sensitivity is not limited to BRCA1 function. Therefore, we asked whether 53BP1 is required for DYNLL1’s role in impeding MRE11 foci formation. To address this question, we fused the FHA domain from RNF8 to the C-terminus of DYNLL1^12^. This strategy has been employed in multiple studies to direct proteins to DSBs circumventing other regulory modules^10, 21^. DYNLL1-EGFP-FHA localized to DNA lesions generated by either irradiation or laser irradiation in BRCA proficient and BRCA1 mutant cell lines in the absence of 53BP1 (Extended Fig. 2C-F). Compared to expression of untethered DYNLL1-EGFP, expression of DYNLL1-EGFP-FHA significantly diminished MRE11 foci after IR-induced DNA damage in BRCA1-deficient cells depleted of 53BP1 or expressing the 53BP1/DYNLL1 binding mutant (Fig. 2D,E and Extended Fig. 2G). Additionally, expression of the FHA-tagged MRE11 binding mutant, DYNLL1^S88A^, failed to suppress MRE11 foci formation after DNA damage (Fig. 2D and Extended Fig. 2G). Together our results suggest that in BRCA1 deficient cells 53BP1 brings DYNLL1 to DSBs to interact with and block MRE11-dependent end resection. However, force tethering DYNLL1 to DSBs ‘rescues’ the DYNLL1-MRE11 interaction, thereby impeding MRE11 foci formation independently of 53BP1. Although this signaling cascade centers around 53BP1, it is distinct from the Shieldin/CST complexes.

**Fig. 2:**
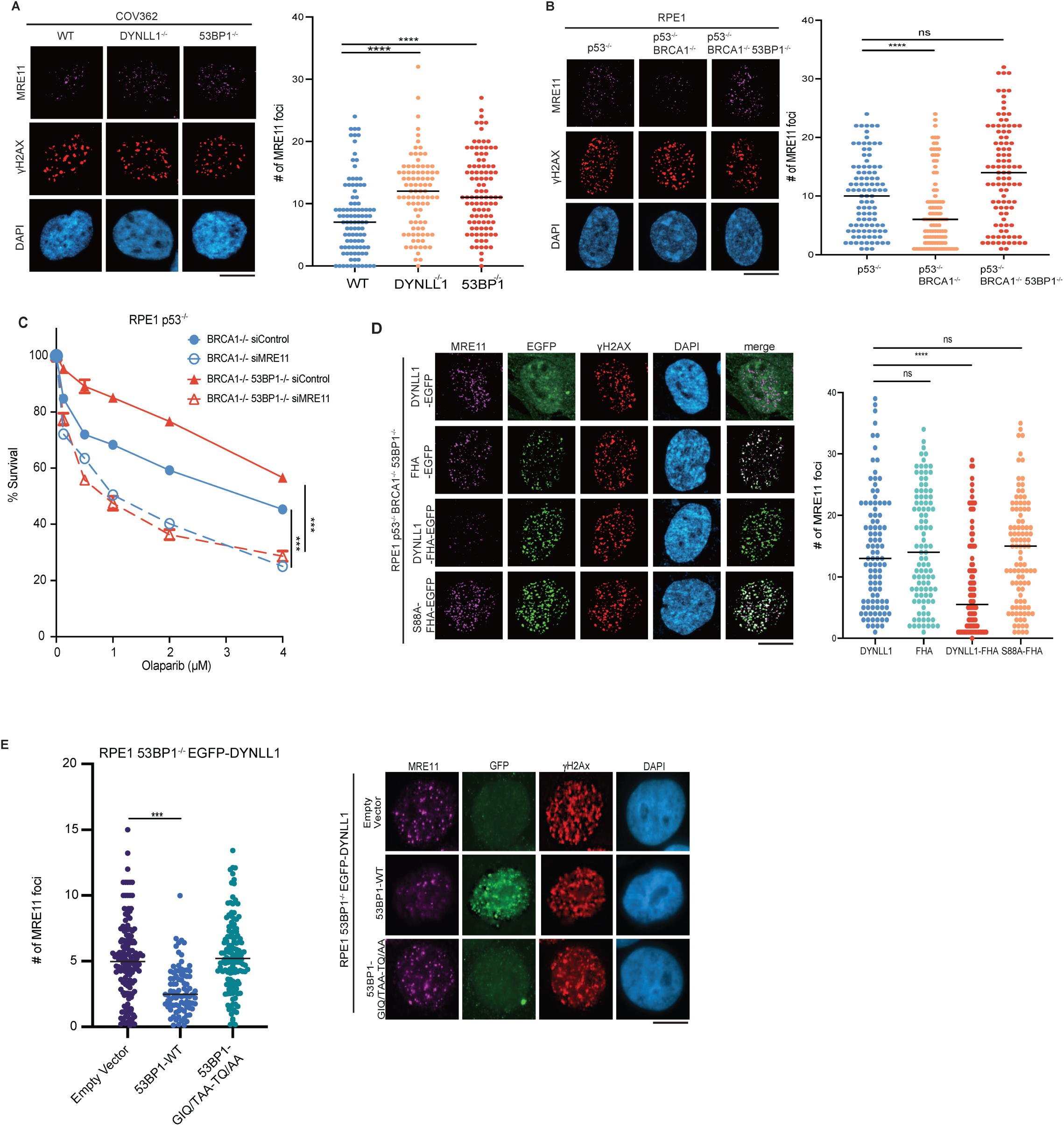
DYNLL1 regulates MRE11 activity independent of 53BP1. (A, B) COV362 cells depleted of DYNLL1 or 53BP1 (A) and RPE1 cells depleted of p53, BRCA1, or 53BP1 (B) using CRISPR/Cas9 were exposed to 2 Gy irradiation. 2 hours post-irradiation cells were fixed and processed for immunofluorescence using antibodies against MRE11 and γH2Ax. (C) RPE1 p53^-/-^ BRCA1^-/-^ and RPE1 p53^-/-^ BRCA1^-/-^ 53BP1^-/-^ cells were depleted of MRE11 using siRNA. Cells were treated with the indicated concentrations of Olaparib for 6 days. Percent survival was determined via a cell viability assay. (D) RPE1 p53^-/-^ BRCA1^-/-^ 53BP1^-/-^ cells expressing GFP-tagged DYNLL1 constructs were exposed to 2 Gy irradiation. 2 hours post-irradiation cells were fixed and processed for immunofluorescence using antibodies against MRE11, GFP (DYNLL1), and γH2Ax. (E) RPE1 53BP1^-/-^ cells coexpressing EGFP-DYNLL1 and 53BP1-DYNLL1 binding mutants were subjected to 2 Gy irradiation and 2 hours later were processed for immunofluorescence using antibodies against MRE11, 53BP1, and γH2Ax. (A-E) For all experiments, n=3 biologically independent experiments, counting ≥ 100 cells per experiment. P-values for foci formation analysis were calculated using the Mann-Whitney test. P-value measurements for cell survival curves were assessed by non-regression curve analysis in Graphpad. Bars on scatter plots represent the median. Error bars represent the mean±s.e.m.

### Functional impact of DYNLL1 at DSBs in 53BP1-deficient cells

RPA directly participates in DSB repair by stimulating 5’-3’ end resection by the BLM helicase and DNA2 endonuclease^22, 23^. To directly test the impact of DYNLL1 on the HR pathway independently of 53BP1, we evaluated RPA foci formation upon expression of DYNLL1-FHA. Expression of FHA-DYNLL1 in BRCA1^-/-^ 53BP1^-/-^ cells reduced the number of RPA foci (Fig. 3A). To directly evaluate the impact of DYNLL1 on end resection we introduced the AsiSI endonuclease fused to estrogen estrogen receptor (ER-AsiSI)^24^ in BRCA1 and 53BP1 depleted cells and used a qPCR-based method^25, 26^ to measure single stranded DNA (ssDNA). Upon expression of DYNLL1-FHA we observed a decrease in ssDNA generated from two specific DSBs (Fig. 3B). Downstream of the DNA end resecting step is loading of RAD51 onto ssDNA, ultimately leading to successful HR mediated DSB repair. BRCA1 loss impairs RAD51 loading and concurrent loss of 53BP1 restores RAD51 foci^6^. Therefore, we evaluated RAD51 foci formation upon expression of DYNLL1-FHA. Expression of FHA-DYNLL1 in BRCA1^-/-^ 53BP1^-/-^ cells reduced the number of RAD51 foci (Fig. 3C and Extended Data Fig.3A-C) and re-sensitized these cells to PARPi (Fig. 3D and Extended Data Fig. 3D) when compared to cells expressing untethered EGFP-DYNLL1. Together these observations suggested that 53BP1 recruits DYNLL1 to DSBs to block MRE11 activity, and this step is sufficient to block HR-mediated DSB repair and sensitize BRCA1-mutant cells to PARPi.

**Fig. 3:**
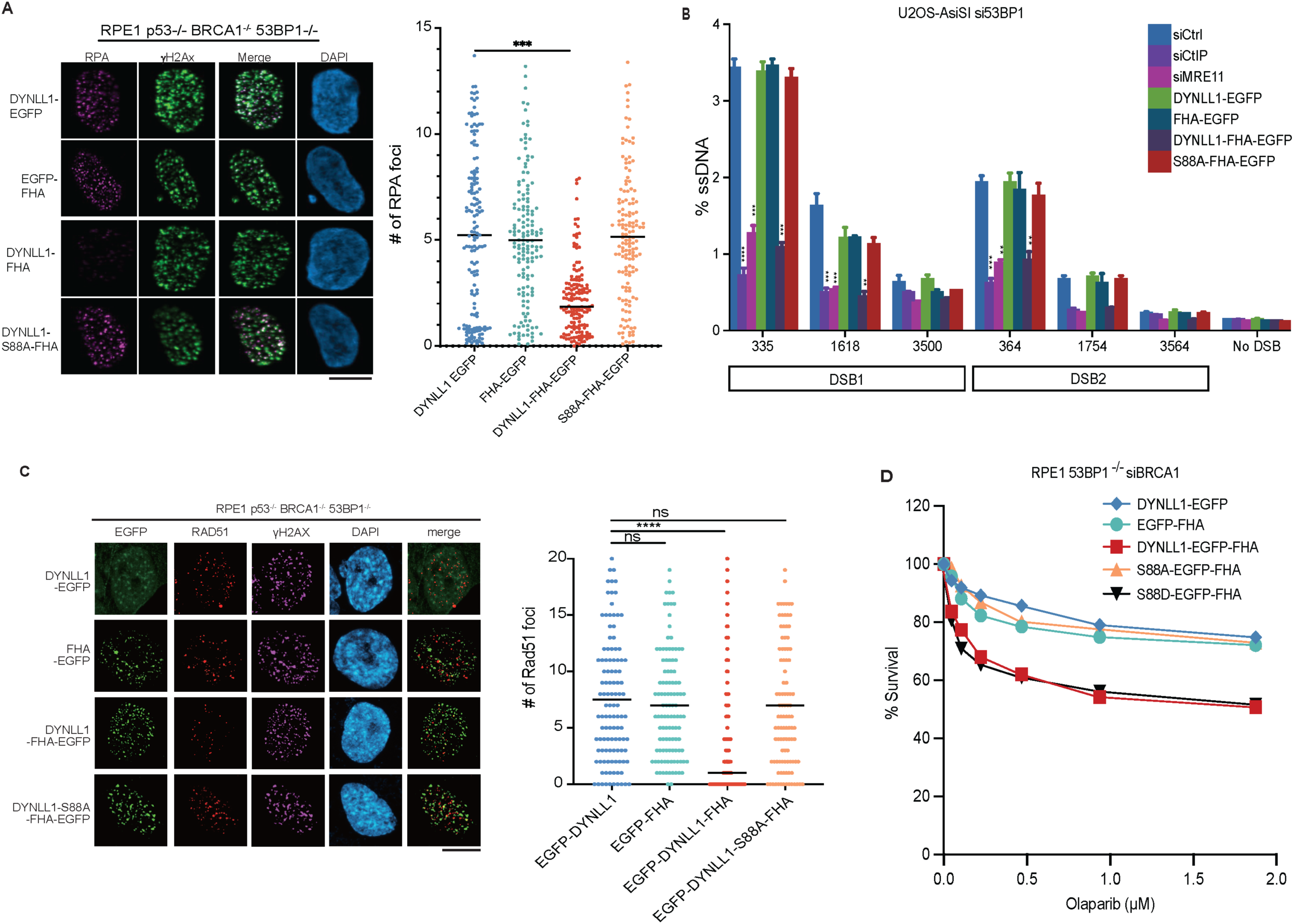
Functional impact of DYNLL1 at DSBs in 53BP1-deficient cells. (A) RPE1 p53^-/-^ BRCA1^-/-^ 53BP1^-/-^ cells expressing GFP-tagged DYNLL or GFP-tagged-FHA-DYNNL1 constructs were exposed to 2 Gy irradiation. 4 hours later, cells were fixed and processed for immunofluorescence using antibodies against GFP (DYNLL1), RPA32, and γH2Ax. (B) U2OS AsiSI cells cotransfected with siRNA against 53BP1 and GFP-DYNLL1 or GFP-FHA-DYNLL1 constructs were treated with 300nM 4-OHT. 4 hours post treatment ssDNA formation at various sites downstream of the break was determined via PCR quantification. (C) RPE1 p53^-/-^ BRCA1^-/-^ 53BP1^-/-^ cells expressing GFP-tagged DYNLL or GFP-tagged-FHA-DYNNL1 constructs were exposed to 2 Gy irradiation. 4 hours later, cells were fixed and processed for immunofluorescence using antibodies against GFP (DYNLL1), RAD51, and γH2Ax. (D) RPE1 p53^-/-^ 53BP1^-/-^ cells were cotransfected with siRNA against BRCA1 and GFP-tagged DYNLL or GFP-tagged FHA-DYNLL1 constructs. Cells were treated with indicated concentrations of Olaparib for 6 days. On day 6, percent survival was determined via a cell viability assay. (A-D) For all experiments, n=3 biologically independent experiments, counting ≥ 100 cells per experiment. P- values for foci quantification were calculated using the Mann-Whitney test. P-values for resection assay were calculated using unpaired t-tests. P-value measurements for cell survival curves were assessed by non-regression curve analysis in Graphpad. Bars on scatter plots represent the median. Error bars represent the mean±s.e.m.

### DYNLL1 disrupts MRE11 dimerization to impair its retention on chromatin

How does DYNLL1 inhibit MRE11 function? To address this question we generated MRE11 truncation mutants and evaluated their interaction with DYNLL1 via immunoprecipitation. (Fig. 4A). Two distinct fragments of MRE11 can bind DYNLL1 in cells. One is the N-terminal fragment (residues 1-181) and the other fragment (residues 293- 483) encompasses a DNA binding domain of MRE11. The N-terminal fragment of MRE11 which includes its dimerization domain is structured while rest of MRE11 is largely disordered (AlphaFold Model-Extended Data Fig. 4A). MRE11 is functionally conserved across species^27^ and crystal structures of MRE11 protein from different species have confirmed that dimerization of MRE11 is evolutionarily conserved^28–31^. Since one of the DYNLL1 binding sites resides in the N-terminus domain of MRE11, we hypothesized that DYNLL1 impacts MRE11 dimerization.

**Fig. 4:**
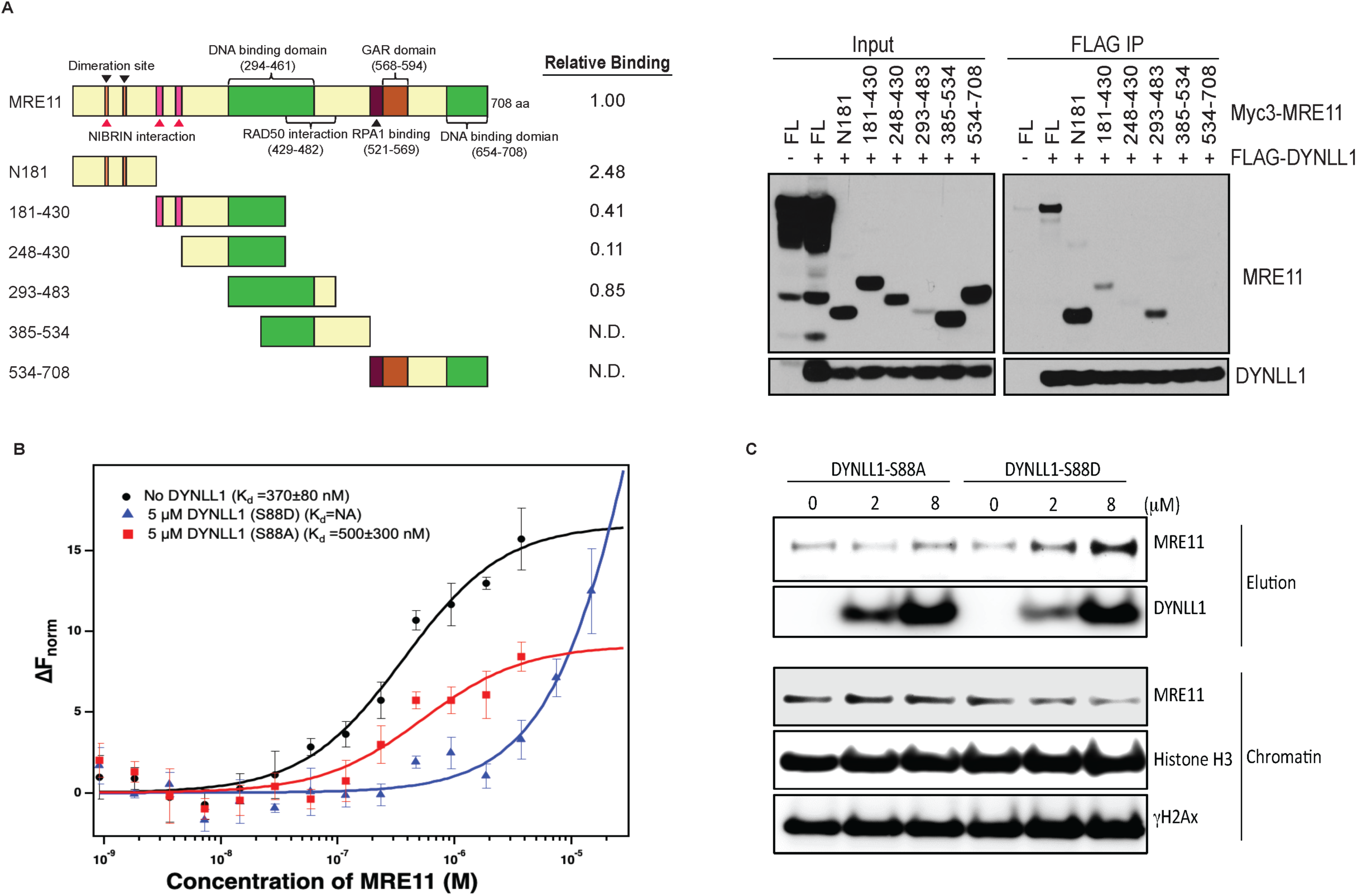
DYNLL1 disrupts MRE11 dimerization to impair its retention on chromatin. (A) Left: Schematic of MRE11 truncation mutants. Right: Relative binding of MRE11 mutants and DYNLL1. Immunoprecipitation from cells coexpressing Flag-DYNLL1 and Myc-MRE11 truncation mutants. N.D.= not detected. (B) Change in the normalized fluorescence as result of thermophoresis in the MST experiment plotted as a function of concentration of unlabeled MRE11. The resulting curves represents MRE11 dimerization in the absence of any DYNLL1 (black circles), in the presence of 5 *μ*M DYNLL1-S88D (blue triangles) or in the presence of 5 *μ*M DYNLL1-S88A (red squares). The K_d_ values are measured by fitting the curves with K_d_ model in the analysis software. The data points represent average of three independent measurements and error bars represents standard deviation. (C) Purified DYNLL1 S88D and DYNLL1 S88A protein (Extended Fig. 4B) was incubated with pre-extracted chromatin from HeLa cells after 5 Gy IR. Western blot shows incubation of recombinant DYNLL1 S88D but not DYNLL1 S88A protein results in increased MRE11 elution from damaged chromatin.

Microscale thermophoresis (MST) can be used to measure the binding constant (Kd) for a protein-protein interaction by titrating unlabeled protein into another labeled protein^32^. Here, we used MST to determine the dimerization Kd of MRE11 by titrating unlabeled MRE11 into Alexa-647-labeled MRE11. In the near-physiological buffer condition we used for the MST experiment, MRE11 forms a dimer with a dimerization Kd of 370±80 nM. To test if DYNLL1 impacts MRE11 dimerization, we repeated MRE11-dimerization MST experiments in the presence of fixed concentration of either DYNLL1-S88D or DYNLL1-S88A mutant in the MST buffer. We observed that MRE11 can still dimerize in the presence of 5 μM DYNLL1-S88A in the MST buffer whereas MRE11 dimerization was almost completely abolished at the same concentration of DYNLL1-S88D (Fig. 4B). This is consistent with our observation that DYNLL1-S88A mutant doesn’t bind MRE11 in cells and S88D binds more efficiently relative to wildtype. We also tested the ten-fold lower concentration of DYNLL1-S88D in the MRE11 dimerization experiment and observed that 500 nM of DYNLL1-S88D is not effective in inhibiting MRE11 dimerization (Extended Data Fig. 4C). Therefore, we concluded that DYNLL1 binds to MRE11 to disrupt its dimerization, which is critical for its function including localization to chromatin.

As a member of the MRN complex, MRE11 is one of the earliest responders to DSBs and is necessary for ATM activation and the expansion of γH2AX foci across megabases surrounding the break site^3, 33, 34^. The 53BP1-dependent recruitment of DYNLL1 to chromatin/DSBs is unlikely to precede MRE11 localization to the DNA repair foci. Therefore, we speculated that DYNLL1 is not preventing the recruitment of MRE11 to DSBs and not interfering with ATM activation. DYNLL1 is more likely to destabilize and remove MRE11 from chromatin by interfering with its dimerization. First we ascertained that DYNLL1 loss doesn’t impact ATM activation (Extended Data Fig. 4D) but leads to increased MRE11 in chromatin^15^. To directly test the impact of DYNLL1 on chromatinized MRE11, we isolated chromatin after IR (5 Gy) from HeLa cells and added recombinant DYNLL1-S88A or DYNLL1-S88D and monitored MRE11 levels. We observed that DYNLL1-S88A had no impact but the addition of DYNLL1-S88D induced a dose-dependent release of MRE11 from chromatin and increase in the soluble fraction (Fig. 4C). Together with our *in vitro* data we infer that DYNLL1 interferes with MRE11 dimerization and promotes its release from chromatin.

### Functional comparison of DYNLL1 and the Shieldin complex

A key question is why cells require multiple and redundant methods to inhibit end resection downstream of 53BP1. Chromosomal aberrations are a readout of the PARPi response^35, 36^. We compared the impact of DYNLL1 and SHLD1 on chromosomal aberrations in the context of PARPi exposure. Depletion of 53BP1, DYNLL1 or SHLD1 in BRCA1 deficient cells had a comparable effect and significantly decreased the number of radials per cell (Fig. 5A). Conversely, expression of DYNLL1-FHA and SHLD1-FHA in BRCA1^-/-^ 53BP1^-/-^ cells increased the number of radials per cell (Fig. 5B), diminished RAD51 foci (Fig. 5C) and resensitized cells to PARPi (Fig. 5D). However, a striking difference was that the expression of DYNLL1-FHA reduced MRE11 foci but SHLD1-FHA had no impact (Fig. 5E). Consistent with these results depletion of SHLD1 also didn’t effect MRE11 foci (Fig. 5F and Extended Data Fig. 5A). These results suggested that the Shieldin complex functions downstream of MRE11 but upstream of Rad51 to impede DNA end resection.

**Fig. 5:**
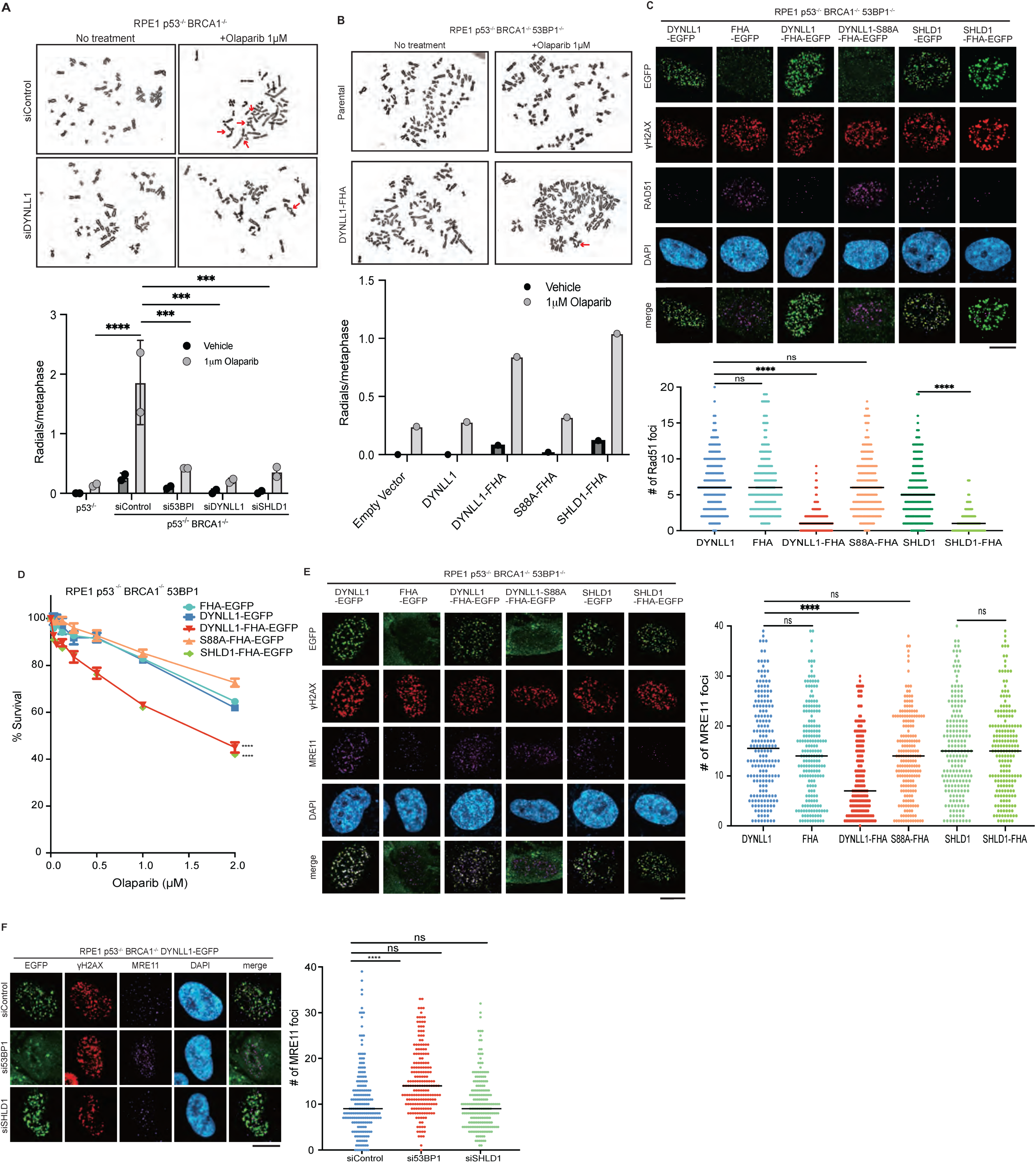
Functional comparison of DYNLL1 and the Shieldin complex. (A) RPE1 p53^-/-^ BRCA1^-/-^ cells depleted of 53BP1, DYNLL1, or SHLD1 were treated with Olaparib. Above: Representative images of metaphase spreads. Below: Quantification of the number of redials/cell. (B) RPE1 p53^-/-^ BRCA1^-/-^ 53BP1^-/-^ cells expressing EGFP-FHA-DYNLL1 constructs treated with Olaparib. Above: Representative images of metaphase spreads. Bottom: Quantification of the number of radials/cell. (C) Cells expressing EGFP-tagged or FHA-tagged DYNLL1, or SHLD1 were exposed to 2 Gy irradiation. 4 hours post-irradiation, cells were fixed and processed for immunofluorescence using antibodies against RAD51. (D) RPE1 p53^-/-^ BRCA1^-/-^ 53BP1^-/-^ cells expressing GFP-tagged DYNLL or GFP-FHA-DYNLL1 and SHLD1 were treated with indicated concentrations of Olaparib. On day 6, percent survival was determined via a cell viability assay. (E) Cells expressing EGFP-tagged or FHA-tagged DYNLL1, or SHLD1 were exposed to 2 Gy irradiation. 2 hours post-irradiation, cells were fixed and processed for immunofluorescence using antibodies against MRE11. (F) RPE1 p53^-/-^ BRCA1^-/-^ 53BP1^-/-^ coexpressing EGFP-DYNLL1 and indicated siRNAs were exposed to 2 Gy irradiation. 2 hours later, cells were fixed and processed for immunofluorescence using antibodies against GFP, γH2Ax, and MRE11. (A-F) Statistics were performed as described in Fig. 2.

### Kinetics and dependencies of Shieldin complex recruitment to DSBs

Next to better understand the dynamics of end resection regulation, we evaluated the kinetics of recruitment of the Shieldin complex relative to 53BP1 and DYNLL1 using laser micro-irradiation and IR. SHLD1 (representative of the Shieldin complex) was recruited to DNA lesions marked by 53BP1, over an hour or more after DYNLL1 (Fig. 6A and Extended Data Fig. 6A, B). When evaluating the kinetics of Shieldin localization to DSBs, we observed that about 20% of 53BP1 foci were occupied by SHLD1 foci, compared to DYNLL1 which occupied about 90% of 53BP1 foci (Fig. 6B), suggesting that Shieldin is recruited to a small number of DSBs and therefore may not participate within the canonical repair pathways.

**Fig. 6:**
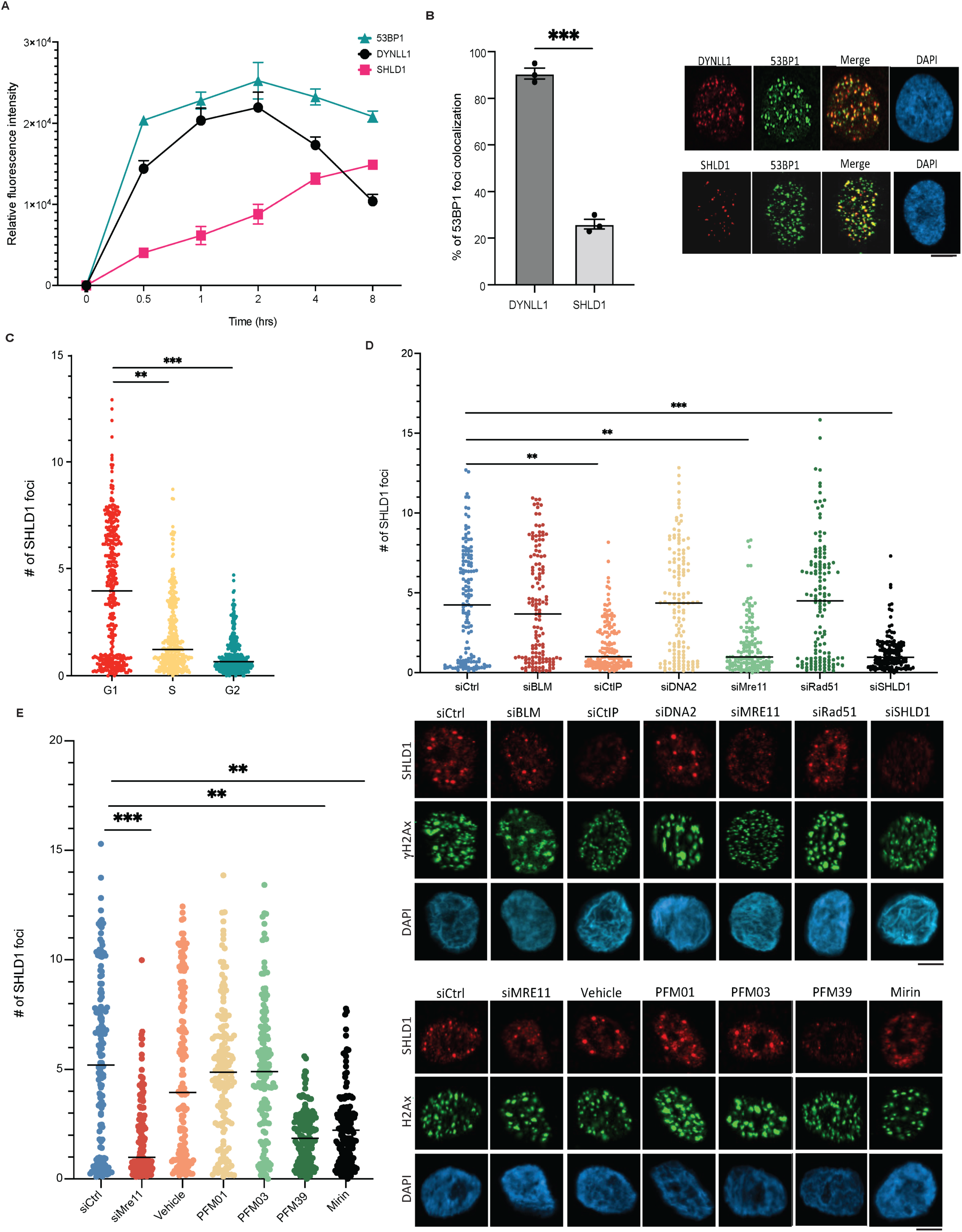
Kinetics and dependencies of Shieldin complex recruitment to DSBs. (A) RPE1 cells were exposed to 10 Gy irradiation and fixed at indicated time points. Cells were processed for immunofluorescence using antibodies against 53BP1, DYNLL1 and γH2Ax. Relative fluorescence intensity is normalized to γH2Ax. (B) RPE1 cells were exposed to 10 Gy irradiation and fixed 4 hours later. Cells were processed for immunofluorescence using antibodies against 53BP1, DYNLL1 and γH2Ax. (C) RPE1 cells were transduced with lentivirus comprised of the Fucci system reporter assay. Cells were exposed to 10 Gy irradiation, fixed 6 hours later, and processed using antibodies against Geminin, Cdt1, and SHLD1. (D, E) RPE1-Fucci cells were transfected with indicated siRNA (D), or treated with MRE11 endo- and exonuclease inhibitors (E). Cells were exposed to 10 Gy irradiation and fixed 6 hours later to be processed for immunofluorescence. Quantification of SHLD1 foci is of cells in G1 phase. (A-E) Statistics were performed as described in Fig. 1.

Next we aimed to determine the cell cycle dynamics of DYNLL1 and Shieldin recruitment to DSBs. We utilized the Fucci cell cycle sensor system, a two-color indicator that uses RFP and GFP fluorescent proteins to follow cell division^37^, and determined that a majority of SHLD1 foci were formed within G1 phase, whereas DYNLL1 recruitment to DSBs was not dependent on cell cycle phase^18^ (Fig. 6C and Extended Data Fig. 6C, D). Given that the Shieldin complex binds to ssDNA via the OB-fold domains of the SHLD2 component^21^, we hypothesized that Shieldin is recruited to ssDNA downstream of end resection in G1. End resection in G1 utilizes specific factors that include MRE11 and CtIP^38^. Depletion of either MRE11 or CtIP, decreased SHLD1 foci formation in G1 phase cells (Fig. 6D and Extended Data Fig. 6E). There was no change in SHLD1 foci formation upon knockdown of G2-dependent end resection factors, BLM and DNA2 (Fig. 6D and Extended Data Fig. 6E). MRE11 endonuclease activity is dispensable for end resection in G1, whereas its exonuclease activity is necessary to promote limited resection at slowly repairing DSBs in G1^38^. Upon inhibition of exonuclease activity, a decrease in SHLD1 foci is observed, whereas cells treated with inhibitors of MRE11’s endonuclease activity didn’t display any affect on SHLD1 localization to DSBs (Fig. 6E). Taken together, we identify that DYNLL1 functions upstream of G1 dependent end resection necessary for Shieldin recruitment in G1 phase cells.

### DYNLL1 is required for Shieldin loading to DSBs

Thus far we have shown that DYNLL1 functions to remove MRE11 from chromatin. Additionally, we have shown that MRE11-dependent minimal end resection is necessary for recruitment of Shieldin to DSBs. Therefore we sought to determine whether DYNLL1 impacts Shieldin recruitment to DSBs. In cells depleted of DYNLL1 we observe an increase of SHDL1 foci at DSBs, and this was negated by expressing DYNLL1-FHA and not by DYNLL1-S88A-FHA (Fig. 7A). Additionally, expression of DYNLL1-S88D decreased SHLD1 foci formation (Fig. 7B) and rescued PARPi sensitivity in BRCA1 and SHLD1 depleted cells (Fig. 7C). This suggests that the interaction of DYNLL1 with MRE11 is critical for SHLD1 recruitment to foci. Notably, cells depleted of 53BP1 but expressing DYNLL1-EGFP or DYNLL1-FHA cannot rescue Shieldin loss (Extended Data Fig. 7A) suggesting that DYNLL1 is not sufficient for Shieldin’s initial localization to DSBs. Furthermore, DYNLL1 does not constitutively interact with any component of the Shieldin complex (data not shown). Together these data suggest that DYNLL1 regulates the recruitment of the Shieldin complex to DSBs via its association with MRE11.

**Fig. 7:**
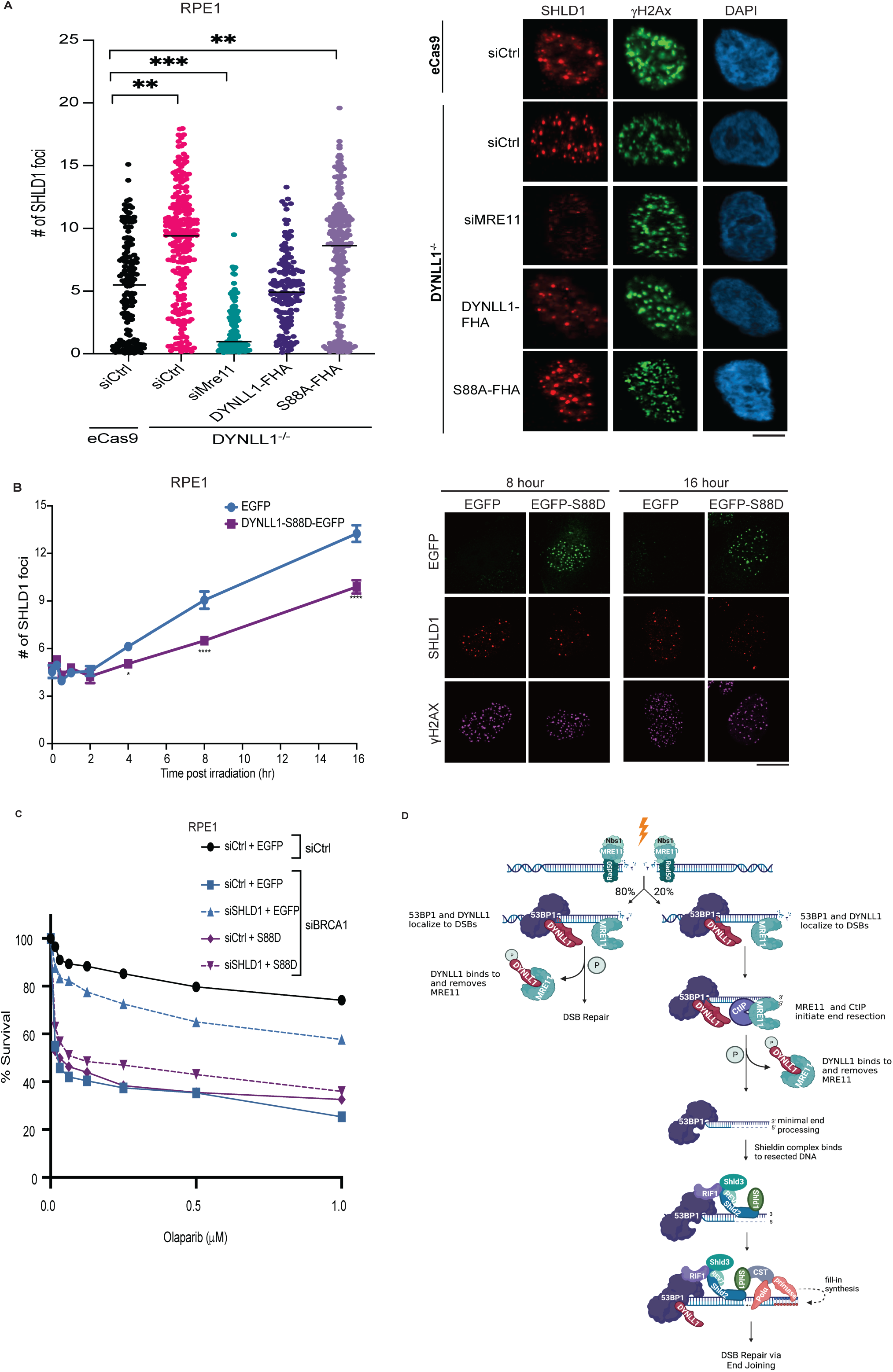
DYNLL1 is required for Shieldin loading to DSBs. (A) RPE1 wild-type or DYNLL1^-/-^ expressing DYNLL1-FHA constructs were exposed to 10 Gy irradiation and fixed 6 hours later for immunofluorescence. (B) RPE1 cells expressing EGFP-DYNLL1 wild-type or its phosphomimetic, S88D were exposed to 10 Gy irradiation. At indicated time points post-irradation, cells were fixed and processed for immunofluorescence using antibodies against GFP (DYNLL1), γH2Ax, SHLD1. (C) BRCA1 depleted RPE1 cells coexpressing DYNLL1 constructs and siRNA against SHLD1 were treated with indicated concentrations of Olaparib. Cell viability assays were performed 6 days after treatment. (D) Model: In about 80% of DSB repair that does not require end resection, phosphorylated DYNLL1-S88 binds to removes MRE11 from chromatin to inhibit resection. In the remaining 20% of repair that occurs during G1, minimal end resection is performed by MRE11 and CtIP. Phosphorylated DYNLL1-S88 then binds to and removes MRE11, allowing for the Shieldin complex to bind to the newly generated ssDNA to protect it from long range end resection by BLM and DNA2.

## DISCUSSION

DNA end resection is a critical step that entails the transient loss of genetic material. In ‘normal’ cells as the resected ssDNA forms a RAD51 nucleofilament and finds the homologous template, the genetic material is restored with *de novo* DNA synthesis. However, the balance of DNA end resection is of paramount importance as excess resection or insufficient resection will impede HR-mediated DSB repair. This equilibrium is often lost in cancer cells as the end resection machinery is harnessed to cause mutagenesis leading to unrestricted proliferation or in other scenarios to induce resistance to DNA damaging cancer therapy. Here we focused on understanding the interplay of the anti-end resection factors centered around the master regulator 53BP1. Our results have defined how distinct 53BP1 interacting proteins work in synchronized chronology to regulate DNA end resection.

DYNLL1 is constitutively associated with 53BP1 and is recruited to DSBs within minutes of damage, and this is dependent on 53BP1 and no other known associated protein. MRE11 is one earliest factors recruited to DSBs with multiple proteins involved in this process^33, 39^. DYNLL1 directly binds MRE11 to remove it from DNA lesions. This step is critical for impeding the initiation of DNA end resection, and potentially promoting NHEJ. In scenarios where MRE11 has been recruited and end resection has occurred, the Shieldin complex is recruited. Therefore, Shieldin complex assembly and recruitment follows DYNLL1 with significant levels of Shieldin proteins visible at DSBs only hours after DNA damage. However, the Shieldin complex recruitment to DSBs does not require DYNLL1 activity, but depends on DYNLL1 being inactive. Phosphorylated DYNLL1 blocks Shieldin recruitment as it impairs the initiation of DNA end resection. Intriguingly, this end resection occurs in a relatively small subset of DSBs (∼20%) predominantly in the G1 phase and is dependent on MRE11 and CtIP. Therefore, Shieldin and CST function downstream of MRE11, and upstream of RAD51. In contrast, DYNLL1 and 53BP1 have roles that are upstream of MRE11 and RAD51. In support of this model (Fig. 7D, Model) we observed that loss of DYNLL1 and 53BP1 enhanced MRE11 and RAD51 foci and forced localization of DYNLL1 at DSBs via the FHA-chimeric protein diminished both MRE11 and RAD51 foci. Tethering SHLD1 blocked only the RAD51 foci and not the MRE11 foci. Although BRCA1 regulates MRE11 function, the impact of MRE11 loss on HR and PARPi sensitivity is beyond BRCA1^40^. This is evident from our observation that MRE11 depletion further sensitizes BRCA1 mutant/deficient cells to PARPi. This further highlights the importance of DYNLL1 mediated regulation of MRE11 activity.

Several questions remain unanswered from these studies. How does the hand-off of DYNLL1 from 53BP1 to MRE11 occur at DSBs? What are the factors involved in this regulatory step? DYNLL1 and Shieldin/CST recruitment to DNA lesions occur independent of one another, although both are reliant on 53BP1. Shieldin complex’s recruitment to damage sites depends on SHLD2’s ability to bind resected ssDNA^7, 12, 41^. However, we need to reconcile how the Shieldin complex inhibits end resection, albeit requiring end resection for its binding to damaged DNA. One possible explanation is that the Shieldin complex inhibits further MRE11-dependent end resection in G1 to re-direct the DSB to NHEJ after fill-in synthesis by the CST complex. In that scenario, we hypothesize (Model, Fig. 7D) that the Shieldin complex is recruited to cleave the ssDNA overhang created by MRE11 end resection using ASTE1 ^42^ and Polα-primase is then recruited for fill-in synthesis and end-joining repair^10, 42^. Another key issue is how the Shieldin complex primarily functioning in G1 influences PARPi sensitivity in BRCA1-mutant tumors. PARPi has been proposed to induce DNA lesions during DNA replication which are then reliant on the BRCA-centric HR pathway. It is feasible that BRCA1 regulates Shieldin proteins and the function of this complex is altered in the context of BRCA1-deficiency. In support of this notion there was increased SHLD1 foci formation in BRCA1-mutant cells^43^. Furthermore, PARPi sensitivity has now been closely tied to replication fork stability and ssDNA gap formation^44^. Loss of REV7^45^ and the CST complex^46^ have been shown to de-stabilize the fork which should cause PARPi sensititivity. This is in contrast to PARPi resistance in BRCA1-mutant cells. Together they suggest that the Shieldin complex may have differential function in absence of BRCA1. Future studies will address these questions and provide further insight on the complex mechanism of DNA end resection and PARPi sensitivity.

## METHODS

### Cell lines and transfection

293T and COV362 cells were cultured in high glucose DMEM (11995065, Gibco) with 10% (v/v) heat-inactivated fetal calf serum (10-437-028, Gibco) and 1% (v/v) Pen-Strep (15140-122, Gibco). RPE1 cells were cultured in DMEM/F12 (10565018, Gibco) with 10% heat-inactivated fetal calf serum and 1% Pen-Strep. U2OS cells were cultured in McCoy’s 5A media with 10% (v/v) heat-inactivated fetal calf serum (10-437-028, Gibco) and 1% (v/v) Pen-Strep (15140-122, Gibco). Cells were maintained in 20% oxygen, 5% CO2, and 37°C. Mouse embryonic fibroblasts (MEFs) were cultured in high glucose and phenol red-free DMEM (21063045, Gibco) with 10% heat-inactivated fetal calf serum. MEF cells were maintained in hypoxic conditions (3% oxygen). All cell lines tested negative for mycoplasma.

All plasmids were transfected using Lipofectamine 2000 (11668019, Invitrogen), and siRNA were transfected by Lipofectamine RNAiMax (13778150, Invitrogen) according to the manufacturer’s protocol.

### Retrovirus and lentivirus production and transduction

HEK293T cells were transfected with various lentiviral or retroviral constructs and viral packaging plasmids. Medium was changed 24 hours after transfection. Virus was collected at 48- and 72-hours post-transfection. All viruses were filtered using a 0.45µm SFCA membrane (723-2545, Thermo Scientific) and used immediately or aliquoted and stored at −80°C.

Virus medium mixed with fresh medium (1:1) was added to 10cm or 6-well plates with 10 µg/mL polybrene (TR-1003, Sigma Aldrich). 24 hours after infection the medium was discarded and changed to fresh medium. Cells expressing GFP or mCherry were selected for by Fluorescence-activated Cell Sorting (FACS).

Western blots showing generation of cell lines is displayed in Supplemental Figure 1.

### Gene knockout and cell line generation

Two CRISPR guide RNAs were selected from GeCKO library^47^. sgRNAs targeting p53, BRCA1, 53BP1, and DYNLL1 were cloned in the pLentiGuide-puro vector (Addgene 52963). 24 hours after viral transduction, cells were selected with puromycin. Three days later, cells were serially diluted, followed by single clonal selection. All knockout cell lines were verified by western blotting.

Western blots showing generation of cell lines is displayed in Supplemental Figure 1.

### Mouse embryonic fibroblasts

Mouse embryonic fibroblasts (MEFs) were generated from E13.5 embryos grown in DMEM supplemented with 15% heat-inactivated fetal bovine serum and 1% pen-strep. Primary MEFs between passages 2-4 were transiently transfected with a vector encoding SV40 T-antigen (pCMV-SV40T) to establish immortalized MEF cell lines. SV40-immortalized MEFs were routinely cultured in DMEM supplemented with 10% FBS.

### Plasmids, antibodies, and reagents

Antibodies and siRNA sequences used in this study can be found in Supplementary Table 1. The siRNAs were used at 20μM. Olaparib (AZD2281) was purchased from Selleckchem (cat# S1060). Concentration and duration of treatment are indicated in the corresponding figures legends.

### Cell Titer-Glo cell viability assay

300-500 cells per well were plated in a 96-well plate. 8 hours later, cells were treated with increasing concentrations of Olaparib (as indicated in corresponding figures) and maintained in drug for six days. The CellTiter-Glo luminescent cell viability assay kit (G9242, Promega) was used to measure cell viability after treatment. The plates were scanned with a luminescence microplate reader. The surviving fraction of drug-treated cells was normalized to values from the DMSO-treated control. Untreated and treated conditions were performed in technical triplicates, and each experiment was repeated at least three times. Survival and statistics were determined by GraphPad Prism software.

### Cell lysis and immunoblotting

For whole-cell protein extraction, cells were harvested using scrapers (CLS3008, Sigma) and first lysed in buffer containing 500mM NaCl, 20 mM HCl [pH 7.5], 0.5% NP-40, 5mM EDTA, 5% glycerol and a protease and phosphatase cocktail inhibitor (Roche). Lysates were rotated at 4°C for 20 minutes. An equal volume of lysis buffer (0mM NaCl, 20 mM HCl [pH 7.5], 0.5% NP-40, 5mM EDTA, 5% glycerol with protease and phosphatase cocktail inhibitor (Roche)) was added, then rotated at 4°C for 20 minutes and spun down at 15,000 rpm for 10 minutes at 4°C. The supernatant was moved to a fresh Eppendorf tube. Protein concentration was measured using the Bradford assay (23225, ThermoFisher Scientific). 50μg of lysate was mixed with SDS loading buffer and loaded into 4%-12% precast gels (NP0335, Life Technologies). For BRCA1 western blotting, 100μg of the lysate was loaded into 3%-8% precast gels (EA0378, Invitrogen™).

For cellular fractionation protein extraction, cells were harvested and lysed in buffer containing 150mM NaCl, 20 mM HCl [pH 7.5], 0.5% NP-40, 5mM EDTA, 5% glycerol and a protease and phosphatase cocktail inhibitor (Roche). Lysates were vortexed to homogenize and incubated on ice for 10 minutes. Lysates were spun down at 2,000 rpm for 4 minutes. The supernatant was transferred to an Eppendorf tube as the soluble fraction. The pellet was gently washed in CE buffer (10mM HEPES, 60mM KCl, 1mM EDTA, 1mM DTT and 1mM PMSF, adjust to pH 7.6) twice. The pellet was resuspended in buffer containing 500mM NaCl, 20 mM HCl [pH 7.5], 0.5% NP-40, 5mM EDTA, 5% glycerol, and a protease and phosphatase cocktail inhibitor (Roche). The pellet was vortexed to homogenize and incubated on ice for 10 minutes. Lysates were spun at 15,000 rpm for 10 minutes. The supernatant was transferred to an Eppendorf tube as the chromatin fraction. For western blotting, 50μg the soluble fraction or 30μg chromatin fraction was mixed with SDS loading buffer and loaded into a precast gel.

### Immunoprecipitation (IP)

Immunoprecipitation was carried out by incubating 1mg of protein lysate with Anti-flag Affinity beads (sigma, A2220) and rotated overnight at 4°C. The following day, the beads were washed and eluted in the SDS loading buffer.

### Immunofluorescence assays

Cells were plated on glass coverslips in 12-well plates. The following day, cells were irradiated at indicated doses and fixed or harvested at indicated time points. Cells were pre-extracted with 0.5% Triton X-100 in CSK buffer (20mM Hepes pH:7.6, 100mM NaCl, 300mM Sucrose, and 3mM MgCl_2_, with a phosphatase inhibitor) for 5 minutes on ice, followed by fixation with 4% paraformaldehyde in CSK buffer for 30 minutes at room temperature. The wells were washed 3 times in PBS containing 3% BSA. Blocking buffer (1%BSA, 10% donkey serum (ab7475, Abcam), 0.1% Triton X-100 in PBS) was added to the plates for 1-hour at room temperature. Cells were incubated with primary antibody overnight at 4°C. Cells were then washed in PBS three times and then incubated with appropriate secondary antibodies for 1 hour at room temperature. Coverslips were rinsed with PBS three times and then mounted using Prolong Gold mounting reagent with DAPI (p36931, Invitrogen). Images were acquired with an Olympus BX41 microscope equipped with a digital camera at 63x magnification.

### Metaphase spreads and chromosomal aberrations

1×10^6^ RPE1 p53^-/-^ BRCA1^-/-^ 53BP1^-/-^ cells were seeded in 10cm plates. 48 hours later cells were treated with 2μM Olaparib for 24 hours. Cells were then treated with colcemid (0.1μg/mL) for 2 hours and harvested using a 0.075M KCl hypotonic solution and fixed with 3:1 methanol:acetic acid. Slides were stained with Wright’s stain, and 50 metaphase spreads were scored for aberrations. The relative number of chromosomal breaks and radials were calculated relative to control cells or empty vector control (indicated in the figure legends).

### Laser microirradiation

Approximately 0.5-1×10^6^ cells were plated in a 35mm μ-Dish with a glass-bottom (81158, Ibidi). On the day of the experiment, the medium was replaced with 1 mL fresh medium. A Zeiss PALM microdissection microscope equipped with a 360nm UV laser at the 38% energy dose was used to create DSBs. All images were analyzed by Fiji (version 2.1.0/1.53c).

### ER-AsiSI resection assay

The percentage of resection adjacent to a specific DSB (DSB1 or DSB2) was measured as previously described^25^. The primer pairs for DSB1 and DSB2 are across BsrG1 and BamHI restriction sites. ER-AsiSI U2OS cells were treated with 4-OHT for 4 hours to allow for the nuclear translocation of AsiSI and induction of DSBs. Cells were harvested and genomic DNA was digested with BsrGI or BamHI enzymes or mock digested overnight at 37°C. Digested samples were used as a template for qPCR. Primers used are listed in the Supplementary Table 1^25^. For each sample, a ΔCt was calculated by subtraction the Ct value of the mock-digested sample from the Ct value of the digested sample.

### Protein expression and purification

N-terminal 6XHis-tagged MRE11 structured-domain (1-411 aa) construct was a kind gift from Dr. Joseph Newman (University of Oxford). This construct was expressed and purified as described previously ^48^ with minor modifications. Briefly, the protein was purified using TALON resin and His-tag was subsequently cleaved with TEV protease during dialysis (in buffer: 25 mM Tris (pH=8), 300 mM NaCl, 2.5% Glycerol, 5 mM B-ME) in the cold room and the tag-cleaved protein was recovered by passing through the TALON^®^ resin again. Protein fractions were concentrated and loaded onto a pre-equilibrated Superdex 200 Increase 10/300 gel-filtration column for further polishing using SEC buffer (20 mM HEPES (pH=7.5), 200 mM NaCl, 1 mM TCEP (pH=7.5)). Both DYNLL1-S88D and DYNLL1-S88A His-tagged recombinant proteins were expressed in chemical competent Rosetta^TM^ cells. DYNLL1 mutant were purified using a similar protocol as in MRE11 using TALON^®^ resin. After His-tag cleavage with TEV protease, untagged proteins were concentrated in the dialysis buffer and flash frozen in liquid nitrogen. Both mutants were further verified by running on a pre-equilibrated Superdex 200 Increase 10/300 gel-filtration column to determine the oligomeric state. As expected, DYNLL1-S88D preferentially forms a monomer whereas the DYNLL1-S88A forms a dimer. Protein quality was further verified by running a protein gel and Coomassie Brilliant Blue G-250 staining.

### Microscale Thermophoresis (MST) to measure the MRE11 dimerization

MRE11 dimerization K_d_ was measured by titrating unlabeled MRE11 into labeled MRE11. MST experiments were performed on a Monolith NT.115Pico system (NanoTemper). Purified MRE11 was fluorescently labelled with an amine-reactive AlexaFluor-647 dye. All MST experiments were performed in the following buffer: 20 mM HEPES (pH=7.5), 150 mM NaCl, 1.5% Glycerol, 0.1 mM EDTA, 1 mM MnCl_2_, 1 mM TCEP and 0.05% T-20. Both ligand (unlabeled MRE11) and target (labeled MRE11) dilution were prepared in the MST buffer. Equal volumes of 10 nM AlexaFluor-647-labelled MRE11 and serially 2X-diluted unlabeled MRE11 were mixed to get a fixed concentration of labelled MRE11 (5 nM) and variable concentration of unlabeled MRE11 (final concentrations ranging from 1.8 nM to 30 μM). The mixture was then incubated for 10 minutes at room temperature before loading into regular Monolith NT.115 capillaries for the MST measurements. After validating the assay condition, the following instrument settings were used for the binding affinity experiments: a 6% excitation power in the Pico-RED channel and high MST power with other default settings including experiment temperature set to 25°C. Each binding experiment was performed in three independent runs and resulting MST data loaded to MO.Analysis (NanoTemper) software to estimate the K_d_ for MRE11 dimerization. To study the effect of DYNLL1 mutants (S88D and S88A) on MRE11 dimerization, MST experiments were performed in the MST buffer with either 5 μM DYNLL1-S88D or 5 μM DYNLL1-S88A. MST data are presented as a change in normalized-fluorescence (due to thermophoresis) as a function of unlabeled MRE11 concentration. The MST data were fit to K_d_ model from the MO.Analysis software. Both raw and fit data were exported and plotted in IgorPro9 software (WaveMetrics) for presentation.

### MRE11 elution from chromatin

Hela cells were treated with 5 Gy IR and pre-extracted with 5% NP-40 lysis buffer with proteinase inhibitor and phosphatase inhibitor. Insoluble chromatin was washed with 1XTBS + 0.05% BSA and incubated with indicated recombinant DYNLL1 protein at 37°C for 30 minutes. Elution was collected and insoluble chromatin was washed with 1X TBS, whereas the chromatin binding protein was extracted with 5% NP-40 lysis buffer + 1% SDS + 0.1 mM EDTA and boiled at 95°C for 10 minutes. Elution and chromatin-bound fractions were subject to Western blot analysis.

### Quantification and statistical analysis

Immunofluorescence foci were counted by CellProfiler (version 4.2.0). Co-localization analyses were done in Fiji (version 2.1.0/1.53c). All statistical analyses were performed by Prism version 8 (GraphPad). All experiments were done in triplicate. P-values for percent “laser positive” analysis was calculated using unpaired t-tests. P-values for foci formation and co-localization were calculated using the Mann-Whitney test. P-value measurements for cell survival curves were assessed by non-regression curve analysis in Graphpad. Error bars represent mean±s.e.m. as described in the figure legends. In all cases, ns: not significant (p≥0.05): *p<0.05, **p<0.01, ***p<0.001, ****p <0.0001.

## DATA AVAILABILITY

All data reported in this paper will be shared by the lead contact upon request. This paper does not report original code.

## ACKNOWLDGEMENTS

D.C. is supported by R01 CA208244 and R01 CA264900, Gray Foundation Team Science Award, DOD Ovarian Cancer Award W81XWH-15-0564/OC140632, Tina’s Wish Foundation, V Foundation Award and the Claudia Adams Barr Program in Innovative Basic Cancer Research. J.A.T. and A.S. were partly supported by NIH grants P01 CA092548, R35 CA220430, plus Cancer Prevention Research Institute of Texas grant RP180813 and a Robert A. Welch Chemistry Chair. We thank Center for Macromolecular Interaction at Harvard Medical School for access to their Monolith NT.115Pico system (NanoTemper). We also thank Dr. Jospeh Newman (University of Oxford) for providing the MRE11 expression construct for producing the recombinant protein.

## AUTHOR INFORMATION

### Contributions

R.Z., M.L.S., and Y.J.H. performed most of the experiments. A.S. purified recombinant proteins and performed MST assays and AlphaFold modeling. K.H. performed immunoprecipitation between DYNLL1 and MRE11 trunction mutants. L.M. performed metaphase spreads and radial formation assay. Y.J.H., J.A.T, A.D.D., P.A.K. and D.C. conceived the study. D.C. wrote the manuscript with contribution from all authors.

### Corresponding authors

Correspondance to Yizhou Joseph He and Dipanjan Chowdhury.

## ETHICS DECLARATIONS

### Competing Interests

The authors declare no competing interests.

## Notes

### Competing Interest Statement

The authors have declared no competing interest.

## REFERENCES

1. Zhao, W., Wiese, C., Kwon, Y., Hromas, R. & Sung, P. The BRCA Tumor Suppressor Network in Chromosome Damage Repair by Homologous Recombination. Annu Rev Biochem 88, 221–245 (2019).

2. Zhao, B., Rothenberg, E., Ramsden, D.A. & Lieber, M.R. The molecular basis and disease relevance of non-homologous DNA end joining. Nat Rev Mol Cell Biol 21, 765–781 (2020).

3. Hustedt, N. & Durocher, D. The control of DNA repair by the cell cycle. Nat Cell Biol 19, 1–9 (2016).

4. Gnugge, R. & Symington, L.S. DNA end resection during homologous recombination. Curr Opin Genet Dev 71, 99–105 (2021).

5. Bunting, S.F. et al. 53BP1 inhibits homologous recombination in Brca1-deficient cells by blocking resection of DNA breaks. Cell 141, 243–54 (2010).

6. Bouwman, P. et al. 53BP1 loss rescues BRCA1 deficiency and is associated with triple-negative and BRCA-mutated breast cancers. Nat Struct Mol Biol 17, 688–95 (2010).

7. Mirman, Z. et al. 53BP1-RIF1-shieldin counteracts DSB resection through CST- and Polalpha-dependent fill-in. Nature (2018).

8. Zimmermann, M. & de Lange, T. 53BP1: pro choice in DNA repair. Trends Cell Biol 24, 108–17 (2014).

9. Ochs, F. et al. 53BP1 fosters fidelity of homology-directed DNA repair. Nat Struct Mol Biol 23, 714–21 (2016).

10. Mirman, Z., Sasi, N.K., King, A., Chapman, J.R. & de Lange, T. 53BP1-shieldin-dependent DSB processing in BRCA1-deficient cells requires CST-Polalpha-primase fill-in synthesis. Nat Cell Biol 24, 51–61 (2022).

11. Mirman, Z. & de Lange, T. 53BP1: a DSB escort. Genes Dev 34, 7–23 (2020).

12. Noordermeer, S.M. et al. The shieldin complex mediates 53BP1-dependent DNA repair. Nature (2018).

13. Dev, H. et al. Shieldin complex promotes DNA end-joining and counters homologous recombination in BRCA1-null cells. Nat Cell Biol (2018).

14. Gupta, R. et al. DNA Repair Network Analysis Reveals Shieldin as a Key Regulator of NHEJ and PARP Inhibitor Sensitivity. Cell (2018).

15. He, Y.J. et al. DYNLL1 binds to MRE11 to limit DNA end resection in BRCA1-deficient cells. Nature 563, 522–526 (2018).

16. Lo, K.W. et al. The 8-kDa dynein light chain binds to p53-binding protein 1 and mediates DNA damage-induced p53 nuclear accumulation. J Biol Chem 280, 8172–9 (2005).

17. West, K.L. et al. LC8/DYNLL1 is a 53BP1 effector and regulates checkpoint activation. Nucleic Acids Res 47, 6236–6249 (2019).

18. Becker, J.R. et al. The ASCIZ-DYNLL1 axis promotes 53BP1-dependent non-homologous end joining and PARP inhibitor sensitivity. Nat Commun 9, 5406 (2018).

19. Setiaputra, D. et al. RIF1 acts in DNA repair through phosphopeptide recognition of 53BP1. Mol Cell 82, 1359–1371 e9 (2022).

20. Mirman, Z. et al. 53BP1-RIF1-shieldin counteracts DSB resection through CST- and Polalpha-dependent fill-in. Nature 560, 112–116 (2018).

21. Noordermeer, S.M. et al. The shieldin complex mediates 53BP1-dependent DNA repair. Nature 560, 117–121 (2018).

22. Chen, H., Lisby, M. & Symington, L.S. RPA coordinates DNA end resection and prevents formation of DNA hairpins. Mol Cell 50, 589–600 (2013).

23. Cejka, P. DNA End Resection: Nucleases Team Up with the Right Partners to Initiate Homologous Recombination. J Biol Chem 290, 22931–8 (2015).

24. Iacovoni, J.S. et al. High-resolution profiling of gammaH2AX around DNA double strand breaks in the mammalian genome. EMBO J 29, 1446–57 (2010).

25. Zhou, Y., Caron, P., Legube, G. & Paull, T.T. Quantitation of DNA double-strand break resection intermediates in human cells. Nucleic Acids Res 42, e19 (2014).

26. Drane, P. et al. TIRR regulates 53BP1 by masking its histone methyl-lysine binding function. Nature 543, 211–216 (2017).

27. Syed, A. & Tainer, J.A. The MRE11-RAD50-NBS1 Complex Conducts the Orchestration of Damage Signaling and Outcomes to Stress in DNA Replication and Repair. Annu Rev Biochem 87, 263–294 (2018).

28. Hopfner, K.P. et al. Structural biochemistry and interaction architecture of the DNA double-strand break repair Mre11 nuclease and Rad50-ATPase. Cell 105, 473–85 (2001).

29. Lammens, K. et al. The Mre11:Rad50 structure shows an ATP-dependent molecular clamp in DNA double-strand break repair. Cell 145, 54–66 (2011).

30. Schiller, C.B. et al. Structure of Mre11-Nbs1 complex yields insights into ataxia-telangiectasia-like disease mutations and DNA damage signaling. Nat Struct Mol Biol 19, 693–700 (2012).

31. Park, Y.B., Chae, J., Kim, Y.C. & Cho, Y. Crystal structure of human Mre11: understanding tumorigenic mutations. Structure 19, 1591–602 (2011).

32. Jerabek-Willemsen, M., Wienken, C.J., Braun, D., Baaske, P. & Duhr, S. Molecular interaction studies using microscale thermophoresis. Assay Drug Dev Technol 9, 342–53 (2011).

33. Paull, T.T. 20 Years of Mre11 Biology: No End in Sight. Mol Cell 71, 419–427 (2018).

34. Paull, T.T. & Lee, J.H. The Mre11/Rad50/Nbs1 complex and its role as a DNA double-strand break sensor for ATM. Cell Cycle 4, 737–40 (2005).

35. de Lange, T. Shelterin-Mediated Telomere Protection. Annu Rev Genet 52, 223–247 (2018).

36. Maciejowski, J. & de Lange, T. Telomeres in cancer: tumour suppression and genome instability. Nat Rev Mol Cell Biol 18, 175–186 (2017).

37. Sakaue-Sawano, A. et al. Visualizing spatiotemporal dynamics of multicellular cell-cycle progression. Cell 132, 487–98 (2008).

38. Biehs, R. et al. DNA Double-Strand Break Resection Occurs during Non-homologous End Joining in G1 but Is Distinct from Resection during Homologous Recombination. Mol Cell 65, 671–684 e5 (2017).

39. Ye, Z. et al. GRB2 enforces homology-directed repair initiation by MRE11. Sci Adv 7(2021).

40. Stracker, T.H. & Petrini, J.H. The MRE11 complex: starting from the ends. Nat Rev Mol Cell Biol 12, 90–103 (2011).

41. Setiaputra, D. & Durocher, D. Shieldin - the protector of DNA ends. EMBO Rep 20(2019).

42. Zhao, F. et al. ASTE1 Promotes Shieldin Complex Mediated DNA Repair by Attenuating End Resection. Nat Cell Biol (2021).

43. Dev, H. et al. Shieldin complex promotes DNA end-joining and counters homologous recombination in BRCA1-null cells. Nat Cell Biol 20, 954–965 (2018).

44. Cantor, S.B. Revisiting the BRCA-pathway through the lens of replication gap suppression: “Gaps determine therapy response in BRCA mutant cancer”. DNA Repair (Amst) 107, 103209 (2021).

45. Paniagua, I. et al. MAD2L2 promotes replication fork protection and recovery in a shieldin-independent and REV3L-dependent manner. Nat Commun 13, 5167 (2022).

46. Lyu, X. et al. Human CST complex protects stalled replication forks by directly blocking MRE11 degradation of nascent-strand DNA. EMBO J 40, e103654 (2021).

47. Sanjana, N.E., Shalem, O. & Zhang, F. Improved vectors and genome-wide libraries for CRISPR screening. Nat Methods 11, 783–784 (2014).

48. Rosenberg, D.J., Syed, A., Tainer, J.A. & Hura, G.L. Monitoring Nuclease Activity by X-Ray Scattering Interferometry Using Gold Nanoparticle-Conjugated DNA. Methods Mol Biol 2444, 183–205 (2022).

