## Supplementary figures and images for "Dynamics of the DYNLL1/MRE11 complex regulates DNA end resection and recruitment of the Shieldin complex to DSBs"

### Extended Figure 1

#### Extended Data Fig. 1

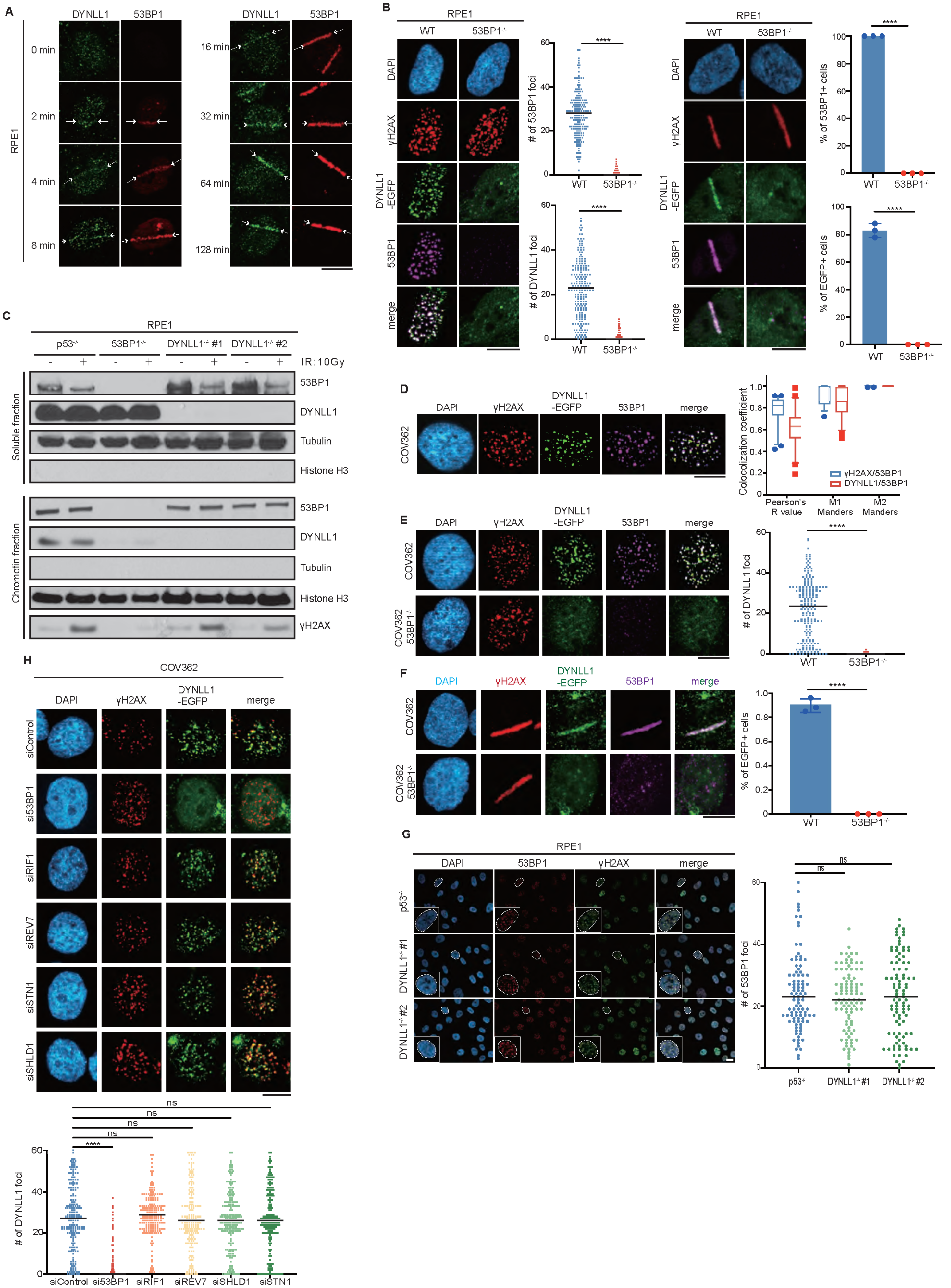

### Extended Figure 2

Extended Data Fig. 2

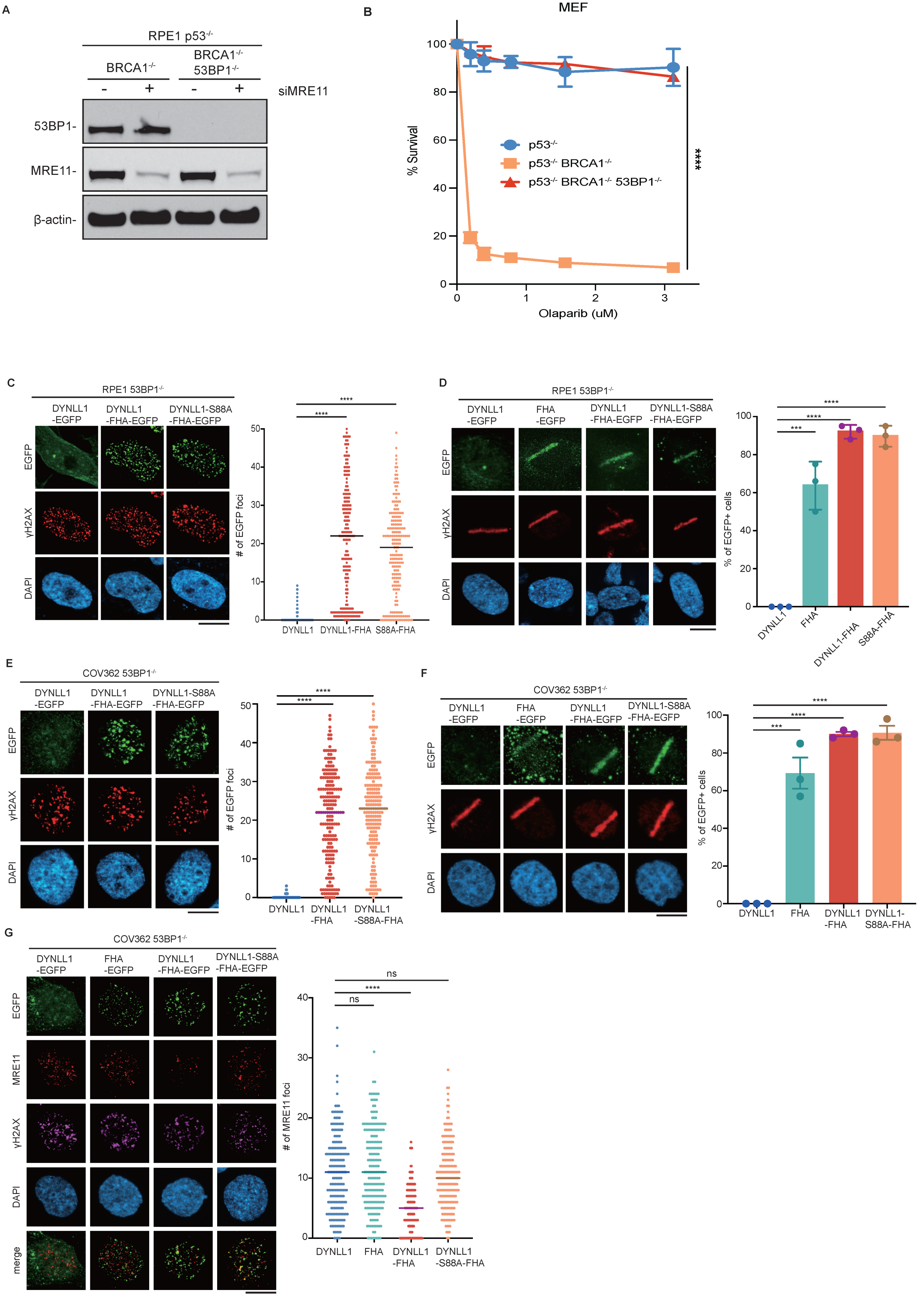

### Extended Figure 3

**Extended Data Fig. 3**

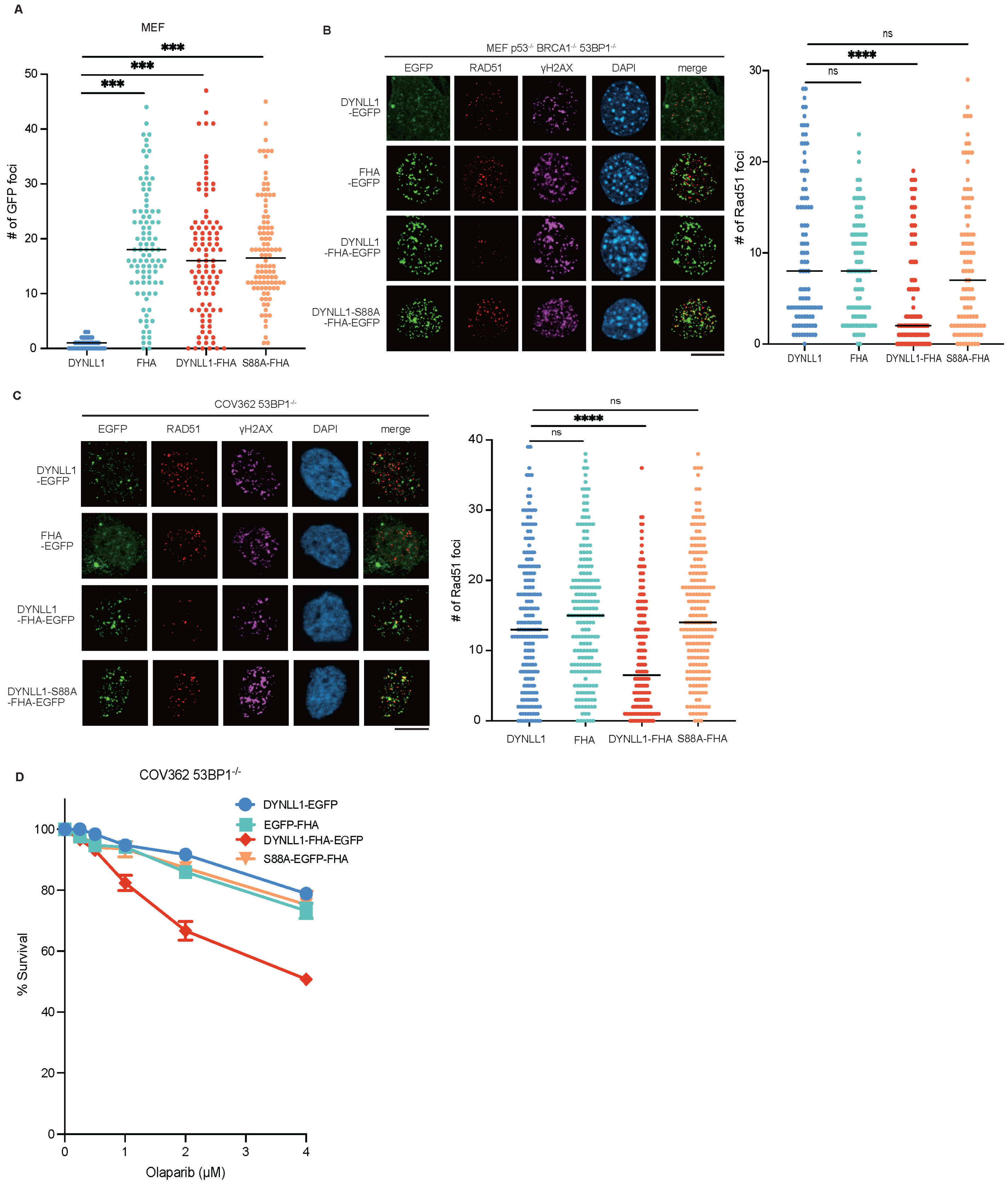

### Extended Figure 4

Extended Data Fig. 4

A

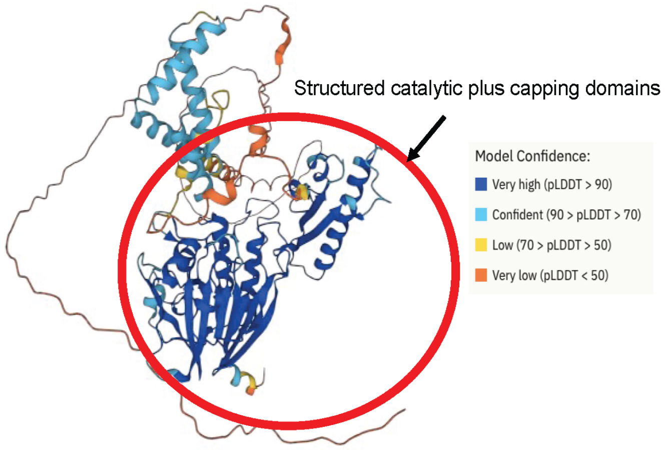


B

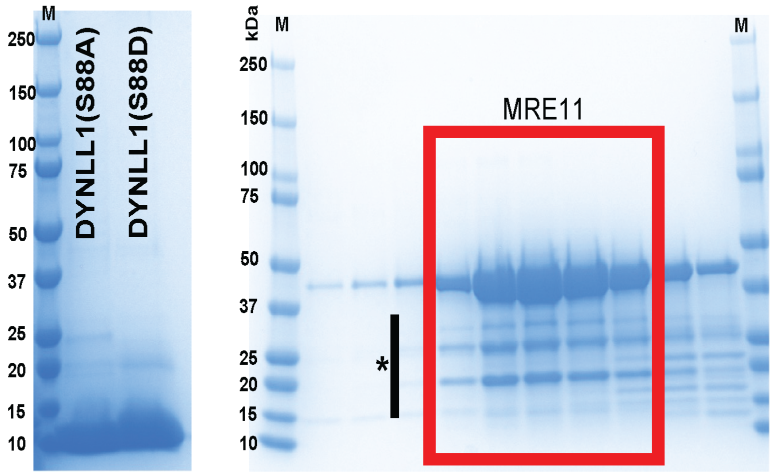


C

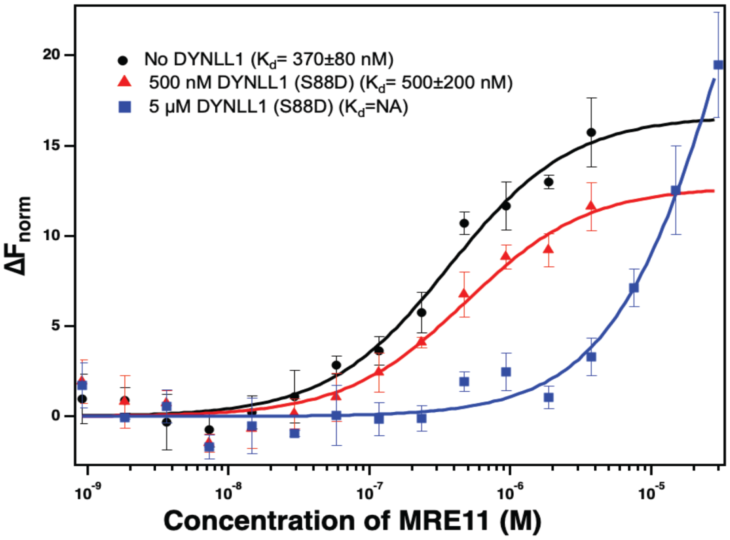


D

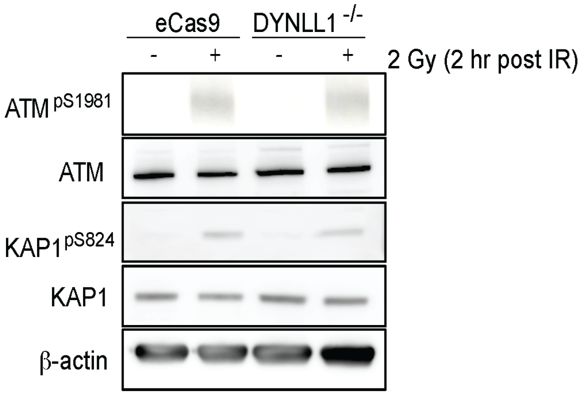

### Extended Figure 5

**Extended Data Fig. 5**

A

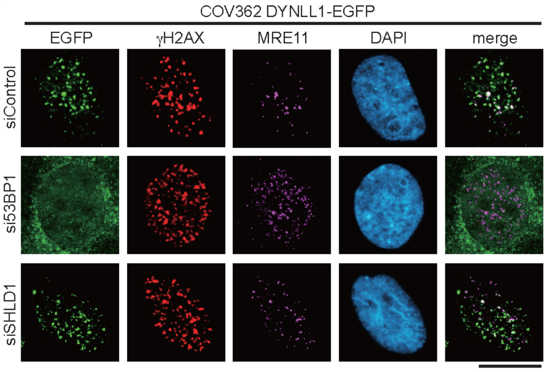


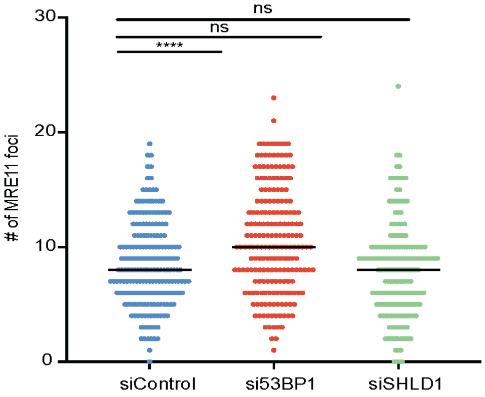

### Extended Figure 6

**Extended Data Fig. 6**

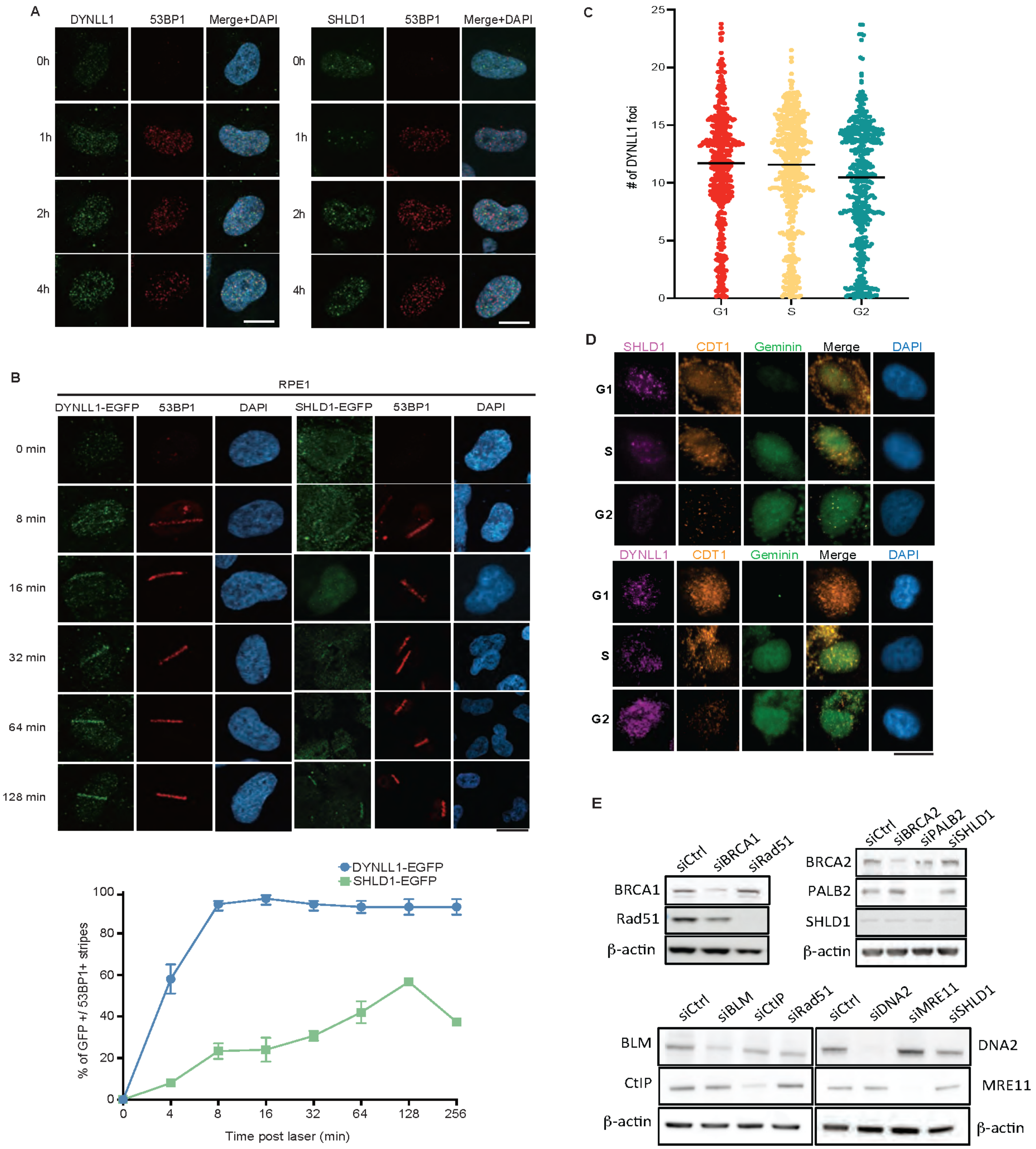

### Extended Figure 7

#### Extended Data Fig. 7

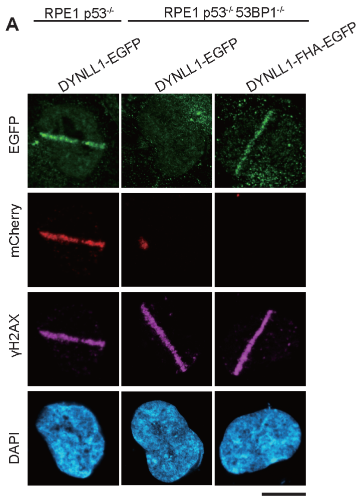

### Supplemental Figure 1

Supplemental Fig. 1

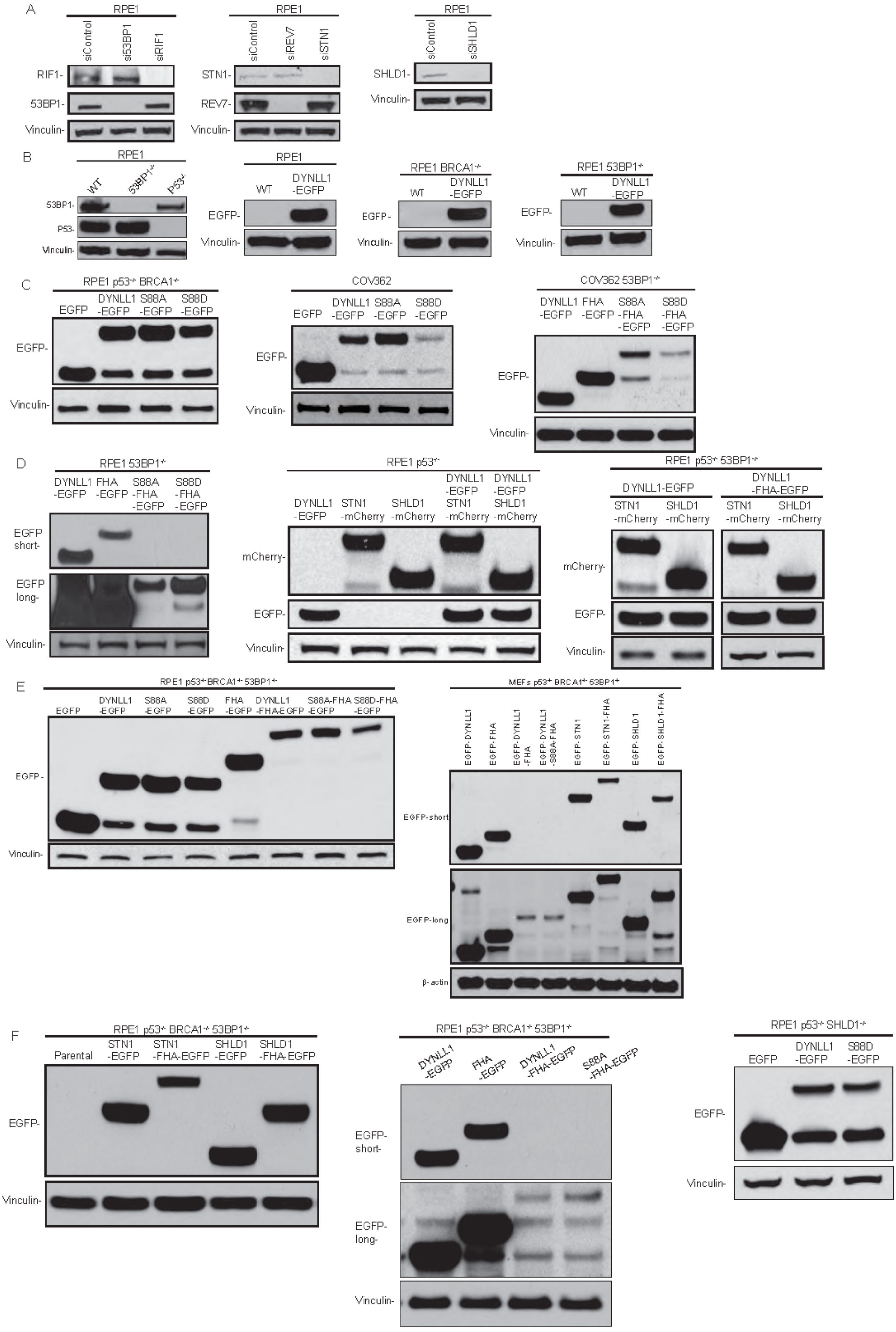
