## Supplementary Table 1 for "Dynamics of the DYNLL1/MRE11 complex regulates DNA end resection and recruitment of the Shieldin complex to DSBs"

| REAGENT or RESOURCE | SOURCE | IDENTIFIER |
|---|---|---|
| **Antibodies** | | |
| GFP (mouse) | Santa Cruz | Sc-9996 |
| GFP (goat) | Abcam | Ab6673 |
| mCherry (mouse) | Novus Biologicals | NBP1-96752 |
| mCherry (rabbit) | Abcam | Ab183628 |
| Rad51 (rabbit) | Abcam | Ab1333534 |
| 53BP1 (rabbit) | Novus Biologicals | NB100-304 |
| 53BP1 (mouse) | Millipore | MAB3802 |
| 53BP1 (mouse) | BD Pharmigen | 612522 |
| γH2Ax (mouse) | Millipore | 05-636 |
| γH2Ax (rabbit) | Cell Signaling Tech. | 9718s |
| γH2Ax (rabbit) | Abcam | Ab11174 |
| Histone H3 (rabbit) | Cell Signaling Tech. | 9715S |
| RIF1 (rabbit) | Bethyl Labs | A300-569A |
| MRE11 (rabbit) | Cell Signaling Tech. | 4895S |
| MRE11 (rabbit) | Novus Biologicals | NB100-143 |
| Tubulin (mouse) | Santa Cruz | Sc-5274 |
| Vinculin (mouse) | Santa Cruz | Sc-25336 |
| B-actin (mouse) | Santa Cruz | Sc-47778 |
| DYNLL1 (rabbit) | Abcam | Ab180579 |
| BRCA1 (mouse) | Millipore | OP92 |
| Myc (rabbit) | Cell Signaling Tech. | 2278S |
| REV7 (rabbit) | Abcam | Ab180579 |
| OBFC1 (STN1) (rabbit) | Abcam | Ab251856 |
| p53 (mouse) | Santa Cruz | Sc-47778 |
| C20orf196 (SHLD1) (rabbit) | Sigma | HPA040749 |
| Flag (mouse) | Sigma | F2555 |
| Flag (mouse) | Sigma | F1804 |
| Cdt1 (goat) | Life Technologies | PA518088 |
| Geminin (mouse) | Abcam | Ab268098 |
| CtIP (rabbit) | Cell Signaling Tech. | 9201S |
| BLM (mouse) | Santa Cruz | Sc-365753 |
| DNA2 (rabbit) | Abcam | Ab96488 |
| RPA32 (rat) | Cell Signaling Tech. | 2208S |
| Donkey anti-Rabbit IgG (H+L) Highly Cross-Adsorbed Secondary Antibody, Alexa Fluor 647 | Invitrogen | A31573 |
| Donkey anti-Mouse IgG (H+L) Highly Cross-Adsorbed Secondary Antibody, Alexa Fluor 568 | Invitrogen | A10037 |
| Donkey anti-Rabbit IgG (H+L) Highly Cross-Adsorbed Secondary Antibody, Alexa Fluor 488 | Invitrogen | A21206 |
| Donkey anti-Goat IgG (H+L) Cross-Adsorbed Secondary Antibody, Alexa Fluor 488 | Invitrogen | A11055 |
| Donkey anti-Rabbit IgG (H+L) Highly Cross-Adsorbed Secondary Antibody, Alexa Fluor 568 | Invitrogen | A10042 |
| Donkey anti-Mouse IgG (H+L) Highly Cross-Adsorbed Secondary Antibody, Alexa Fluor 647 | Invitrogen | A31571 |
| HRP Secondary Rabbit antibody | Jackson Immuno Res. | 711-035-152 |
| HRP Secondary Mouse antibody | Jackson Immuno Res. | 715-035-150 |

### Supplementary Table 1

| | | |
|---|---|---|
| **Experimental models: Cell lines** | | |
| 293T | ATCC | CRL-3216 |
| COV362 | Sigma | 07071910 |
| hTert-RPE1 | ATCC | CRL-4000 |
| U2OS-AsiSI | Gaelle Legube | |
| Mouse embryonic fibroblasts (MEF) | DFCI | |
| **Oligonucleotides** | | |
| siControl: 5’-AAGCCGGUAUGCCGGUUAAGU-3′ | Dharmacon | |
| si53BP1:5‘-AGAACGAGGAGACGGUAAUAGUGGG-3' | Dharmacon | |
| siMRE11_1: 5‘-CGAAAUGUCACUAAGAUU-3' | Dharmacon | |
| siMRE11_2: 5‘-GGAGGUACGUCGUUUCAGAUU-3' | Dharmacon | |
| siREV7: 5‘-AAGAUGCAGCUUUACGUGGAA-3'; | Dharmacon | |
| siSTN1: 5‘-GCUUAACCUCACAACUUAA-3' | Dharmacon | |
| siBRCA1: 5′-CAGCUACCCUUCCAUCAUA-3′ | Dharmacon | |
| siDYNLL1: 5′-GAAGGACAUUGCGGCUCAU-3′ | Dharmacon | |
| siRIF1: 5‘-GACUCACAUUUCCAGUCAA-3' | Dharmacon | |
| siSHLD1: 5‘-CAGCGAGGCUUUCAGUUCU-3' | Dharmacon | |
| siBLM: 5’-AAGCTAGGAGTCTGCGTGCGA-3’ | Dharmacon | Biehs et al, Mol Cell. 2017 |
| siDNA2: 5’-AAATAGCCAGTAGTATTCGAT-3’ | Dharmacon | Biehs et al, Mol Cell. 2017 |
| siCtIP: 5’-TCCACAACATAATCCTAATAA-3’ | Dharmacon | Biehs et al, Mol Cell. 2017 |
| **Primers for ER-AsiSI** | | |
| DSB1-335bp_F: GAATCGGATGTATGCGACTGATC<br>DSB1-335bp_R: TTCCAAAGTTATTCCAACCCGAT | Dharmacon | Zhou et al, Nuc. Acids Res. 2014 |
| DSB1-1618bp_F: TGAGGAGGTGACATTAGAACTCAGA<br>DSB1-1618bp_R: AGGACTCACTTACACGGCCTTT | Dharmacon | Zhou et al, Nuc. Acids Res. 2014 |
| DSB1-3500_F: TCCTAGCCAGATAATAATAGCTATACAAACA<br>DSB1-3500_R:TGAATAGACAGACAACAGATAAATGAGACA | Dharmacon | Zhou et al, Nuc. Acids Res. 2014 |
| DSB2-364_F: CCAGCAGTAAAGGGGAGACAGA<br>DSB2-364_R: CTGTTCAATCGTCTGCCCTTC | Dharmacon | Zhou et al, Nuc. Acids Res. 2014 |
| DSB2-1754bp_F: GAAGCCATCCTACTCTTCTCACCT<br>DSB2-1754bp_R: GCTGGAGATGATGAAGCCCA | Dharmacon | Zhou et al, Nuc. Acids Res. 2014 |
| DSB2-3564bp_F: GCCCAGCTAAGATCTTCCTTCA<br>DSB2-3564bp_R: CTCCTTTGCCCTGAGAAGTGA | Dharmacon | Zhou et al, Nuc. Acids Res. 2014 |
| No DSB_F: ATTGGGTATCTGCGTCTAGTGAGG<br>No DSB_R: GACTCAATTACATCCCTGCAGCT | Dharmacon | Zhou et al, Nuc. Acids Res. 2014 |
| Across DSB1_F: GATGTGGCCAGGGATTGG<br>Across DSB1_R: CACTCAAGCCCAACCCGT | Dharmacon | Zhou et al, Nuc. Acids Res. 2014 |
| Across DSB2_F: GAGGAGCCTCTCCTGCAGC<br>Across DSB2_R: GAACCAGACCTACCTCCAGGG | Dharmacon | Zhou et al, Nuc. Acids Res. 2014 |
| **Recombinant DNA** | | |
| pBOB-EF1-FastFUCCI-Puro | Addgene | 86849 |
