## Extended and Supplementary Figure Legends for "Dynamics of the DYNLL1/MRE11 complex regulates DNA end resection and recruitment of the Shieldin complex to DSBs"

**Extended Data Fig. 1: 53BP1 is necessary for chromatin localization of DYNLL1.**

**Related to Fig 1.**

(A) Representative immunofluorescent images of RPE1 cells subjected to laser microirradiation. Cells were fixed at indicated time points post laser microirradiation and processed for immunofluorescence with DYNLL1 and 53BP1 antibodies. (B) Representative images of RPE1 wild-type or 53BP1$^{-/-}$ cells 2 hours after exposure to 2 Gy irradiation or laser microirradiation. Cells were fixed and processed for immunofluorescence using antibodies against 53BP1, GFP (DYNLL1), and γH2Ax. (C) RPE1 cells depleted of p53, 53BP1, or DYNLL1 using CRISPR/Cas9 were exposed to 10 Gy irradiation. Protein was collected after 3 hours. Localization of DYNLL1 to chromatin was evaluated by subcellular fractionation followed by immunoblotting for DYNLL1. (D-F) Representative images of COV362 cells (D) and COV362 wild-type or 53BP1$^{-/-}$ cells (E, F) exposed to 2 Gy irradiation (D, E) or laser microirradiation (F). 2 hours post-recovery cells were fixed and processed for immunofluorescence using antibodies against 53BP1, GFP (DYNLL1), and γH2Ax. (G) Representative immunofluorescent images of RPE1 cells depleted of DYNLL1 using CRISPR/Cas9 and exposed to 2 Gy irradiation. 2 hours post-irradiation, cells were fixed and processed for immunofluorescence using antibodies against 53BP1 and γH2Ax. (A-F) Statistics were performed as described in Fig. 1.

**Extended Data Fig. 2: Force tethering DYNLL1 to chromatin inhibits MRE11 foci formation.**

**Related to Fig. 2.**

(A) Representative western blot for Fig. 2C. (B) MEF cells depleted of p53, or p53 and BRCA1, or p53, BRCA1 and 53BP1 were treated with indicated doses of Olaparib for 6 days. Percent survival was determined by a cell viability assay. (C, D) RPE1 53BP1$^{-/-}$ cells were transfected with GFP-tagged DYNLL1 or DYNLL1-FHA constructs. Cells were subjected to 2 Gy irradiation (C) or laser microirradiation (D). 2 hours later cells were fixed and processed for immunofluorescence using antibodies against GFP (DYNLL1) and γH2Ax. (E, F) COV362 53BP1$^{-/-}$ cells were transfected with a GFP-tagged DYNLL1 or DYNLL1-FHA constructs. Cells were subjected to 2 Gy irradiation (E) or laser microirradiation (F). 2 hours later cells were fixed and processed for immunofluorescence using antibodies against GFP (DYNLL1) and γH2Ax. (G) COV362 53BP1$^{-/-}$ cells were exposed to 2 Gy irradiation. 2 hours post-irradiation cells were fixed and processed for immunofluorescence using antibodies against MRE11 and γH2Ax. (A-G) Statistics performed as described in Fig. 2.

**Extended Data Fig. 3: DYNLL1 chromatin binding suppresses 53BP1 loss-induced restoration of HR in BRCA1 deficient cells.**

**Related to Fig. 3.**

(A) MEFs expressing GFP-tagged DYNLL1, and GFP-tagged DYNLL1-FHA domains constructs were exposed to 2 Gy irradiation. 2 hours after irradiation cells were fixed and processed for immunofluorescence using a GFP (DYNLL1) antibody. (B, C) MEF p53$^{-/-}$ BRCA1$^{-/-}$ 53BP1$^{-/-}$ cells (B) and COV362 53BP1$^{-/-}$ cells (C) were transfected with a GFP-tagged DYNLL, or GFP-tagged DYNNL1-FHA constructs. Cells were exposed to 2 Gy irradiation. 2 hours later, cells were fixed and processed for immunofluorescence using antibodies against GFP (DYNLL1), RAD51, and γH2Ax. (D) COV362 53BP1$^{-/-}$ cells were transfected GFP-tagged DYNLL, or GFP-tagged DYNNL1-FHA constructs. Cells were treated with indicated concentrations of Olaparib for 6 days. Percent survival was determined via a cell viability assay. (A-D) Statistics performed as described in Fig. 2.

**Extended Data Fig. 4: DYNLL1 interferes with MRE11 dimerization.**

**Related to Fig. 4.**

(A) Predicted structure of full-length MRE11 created by AlphaFold Monomer V2 for Uniprot Accession number P49959. The structured catalytic domain of MRE11 is highlighted with a red circle. The model is color coded in terms of confidence in prediction and respective color schemes for the confidence is given in the figure. In general, disordered regions have less confidence in model prediction, thus indicating the unstructured regions of MRE11 beyond capping domain. (B) Coomassie-stained protein gels indicating the quality of the recombinant protein used in the current study. Left: DYNLL1 mutants after cleaving the His-tag with TEV protease. Right: MRE11 catalytic domain after the gel-filteration purification step. The red rectangle indicates the fractions that are combined. M indicates the protein standards and * indicates the MRE11 degradation bands. (C) Change in the normalized fluorescence as result of thermophoresis in the MST experiment plotted as a function of concentration of unlabeled MRE11. The resulting curves represents MRE11 dimerization in the absence of any DYNLL1 (black circles), in the presence of 500 nM DYNLL1-S88D (red triangles) or in the presence of 5 $\mu$M DYNLL1-S88D (blue squares). The $K_d$ values are measured by fitting the curves with $K_d$ model in the analysis software. The data points represent average of three independent measurements and error bars represents standard deviation. (D) Protein expression from lysates collected from RPE1 wild-type or DYNLL1$^{-/-}$ cells 2 hours after 2 Gy irradiation.

**Extended Data Fig. 5: Depletion of the Shieldin complex does not affect MRE11 recruitment. Related to Fig. 5.**

(A) COV362 cells overexpressing EGFP-DYNLL1 and transfected with siRNA targeting either 53BP1, or SHLD1 were exposed to 2 Gy irradiation. Cells were fixed 2 hours post exposure and processed for immunofluorescence using antibodies against GFP (DYNLL1), γH2Ax, and MRE11. Statistics were performed as described in Fig. 1.

**Extended Data Fig. 6: Shieldin is recruited to DSBs later than DYNLL1 and in G1 phase only. Related to Fig. 6.**

(A) Representative images from Fig. 6A. (B) Representative images from cells expressing DYNLL1-EGFP or SHLD1-EGFP and subjected to laser microirradiation. Cells were fixed at indicated time points and processed for immunofluorescence. (C) RPE1 cells were transduced with lentivirus comprised of the Fucci system reporter assay. Cells were exposed to 10 Gy irradiation, fixed 6 hours later, and processed using antibodies against Geminin, Cdt1, and DYNLL1. (D) Representative images for Fig. 6C and S6C. (E) Representative western blots showing knockdown of indicated proteins for Fig. 6D. (A-C) Statistics were performed as described in Fig. 1.

**Extended Data Fig. 7: Shieldin functions downstream of DYNLL1, but is not dependent on DYNLL1 for its localization to chromatin.**

**Related to Fig. 7.**

(A) RPE1 p53$^{-/-}$ and RPE1 p53$^{-/-}$ 53BP1$^{-/-}$ cells were transfected with GFP-tagged DYNLL1 or GFP-tagged DYNLL1-FHA. Cells were then transfected with mCherry-SHLD1 and subjected to laser microirradiation. 2 hours after laser microirradiation cells were fixed and processed for immunofluorescence using antibodies against GFP (DYNLL1), mCherry (SHLD1), and γH2Ax. Statistics were performed as described in Fig. 1.

**Supplementary Fig. 1**

Contains western blots of generated cell lines.

**Supplementary Table 1**

Lists key resources, including chemical reagents, cell lines, antibodies, primer sequences, and plasmids used within the study.
